# Cortical and subcortical hemodynamic changes during human sleep slow waves

**DOI:** 10.1101/2020.04.29.067702

**Authors:** Monica Betta, Giacomo Handjaras, Andrea Leo, Alessandra Federici, Valentina Farinelli, Emiliano Ricciardi, Francesca Siclari, Stefano Meletti, Daniela Ballotta, Francesca Benuzzi, Giulio Bernardi

## Abstract

EEG slow waves, the hallmarks of NREM sleep, are closely linked to the restorative function of sleep and their regional cortical distribution reflects plasticity- and learning-related processes. Here we took advantage of simultaneous EEG-fMRI recordings to map cortical and subcortical hemodynamic (BOLD) fluctuations time-locked to sleep slow waves. Recordings were performed in twenty healthy adults during an afternoon nap. Slow waves were associated with BOLD-signal increases in the brainstem and in portions of thalamus and cerebellum characterized by preferential functional connectivity with limbic and somatomotor areas, respectively. At the cortical level, significant BOLD-signal decreases were found in several areas, including insula and somatomotor cortex, and were preceded by slow signal increases that peaked around slow-wave onset. EEG slow waves and BOLD fluctuations showed similar cortical propagation patterns, from centro-frontal to temporo-occipital cortices. These regional patterns of hemodynamic-electrical coupling are consistent with theoretical accounts of the functions of sleep slow waves.

## Introduction

Slow waves of sleep are thought to be crucial for the regulation of several important sleep-related processes, including sensory disconnection and synaptic plasticity related to learning and memory consolidation (Crunelli et al., 2018; Timofeev and Chauvette, 2017; Tononi and Cirelli, 2014). Recent work also suggested that a direct relationship may exist between electroencephalographic (EEG) slow waves, hemodynamic and cerebrospinal fluid (CSF) dynamics, ultimately resulting in the removal of metabolic wastes from the brain (Fultz et al., 2019; Hablitz et al., 2019; Xie et al., 2013).

The appearance of NREM slow waves in the EEG signal depends on the coordinated oscillations of cortical neuronal populations between a hyperpolarized *down-state* with neuronal silence and a depolarized *up-state* characterized by intense neuronal firing (Steriade et al., 1993). While this *slow oscillation* appears in both cortical and thalamic networks, its origin is believed to be cortical. In fact, slow waves persist in the neocortex after thalamic lesions or pharmacological blockade of thalamic activity (David et al., 2013; Steriade et al., 1993). Moreover, slow waves are expressed in isolated cortical slabs *in vivo* and in cortical slice preparations *in vitro* (Crunelli and Hughes, 2010; Lőrincz et al., 2015; Sanchez-Vives and McCormick, 2000; Timofeev et al., 2000). On the other hand, the *slow oscillation* disappears in the thalamus of decorticated animals (Timofeev and Steriade, 1996). However, a growing body of evidence indicates that the thalamus and other subcortical structures, including the basal forebrain and several brainstem nuclei, may have an active role in regulating the expression of cortical slow waves in physiological conditions (Gent et al., 2018b; Neske, 2016). Indeed, an increase in thalamic activity has been shown to precede the initiation of cortical *up-states* in animal models (Gent et al., 2018a; Sheroziya and Timofeev, 2014; Slézia et al., 2011; Ushimaru and Kawaguchi, 2015). Moreover, thalamic deafferentation of the cortex results in a reduced frequency of the *slow oscillation* (David et al., 2013; Lemieux et al., 2014).

At the cortical level, each slow wave behaves as a traveling wave characterized by a specific origin and propagation pattern (Massimini et al., 2004; Menicucci et al., 2009; Murphy et al., 2009). Interestingly, slow waves appear to originate more often within the somatomotor cortex and the insula (Massimini et al., 2004; Murphy et al., 2009), from which they spread towards anterior and posterior brain areas, especially along the medial surfaces of the two brain hemispheres (Menicucci et al., 2009; Murphy et al., 2009). Such a propagation seems to mostly occur through cortico-cortical white matter connections (Avvenuti et al., 2020; Buchmann et al., 2011; Murphy et al., 2009; Piantoni et al., 2013). Importantly, the organized and systematic traveling of slow waves along connected pathways has been suggested to have an important role in sleep-dependent network-level processes related to plasticity and memory consolidation (Cox et al., 2014; Massimini et al., 2004).

In summary, slow waves of sleep appear to involve complex interactions among multiple cortical and subcortical structures. Understanding such interactions is crucial to improve our knowledge regarding the regulation and functional role of (NREM) sleep, as well as its alterations under pathological conditions. However, conventional EEG investigations in humans have a low spatial resolution and are unable to accurately describe changes in the activity of subcortical and deep cortical structures. These issues may be overcome by combining the optimal temporal resolution of EEG with the high spatial resolution offered by functional magnetic resonance imaging (fMRI; Mullinger and Bowtell, 2010). In fact, one EEG-fMRI investigation on slow-wave correlates reported significant BOLD-signal increases in several brain areas, including the brainstem, cerebellum, inferior frontal cortex, precuneus and posterior cingulate areas (Dang-Vu et al., 2008). Surprisingly, however, no clear changes in thalamic activity were found in association with sleep slow waves. Moreover no cortical regions showed significant decreases in brain activity despite the well-known association of slow waves with a highly synchronized suppression of neuronal firing (Steriade et al., 1993). These inconsistencies could have their roots in the non-stationary nature of sleep slow waves, as conventional analyses may fail to capture time-varying changes in brain activity (Mitra et al., 2015). Furthermore, the previous investigation only focused on large-amplitude slow waves and was thus unable to determine how slow-wave amplitude relates to changes in hemodynamic activity measured through fMRI. In order to address these issues, here we took advantage of simultaneous EEG-fMRI recordings to obtain spatially and temporally detailed maps of cortical and subcortical hemodynamic fluctuations time-locked to sleep slow waves (0.5-2 Hz).

## Results

### Final sample and sleep macrostructure

Simultaneous EEG-fMRI recordings were performed in twenty healthy adults during an afternoon nap. However, three subjects (all males) were excluded from further evaluation as they did not reach stable sleep (n=1) or presented strong artifactual activity in the EEG-signal (n=2). Thus, the final sample included 17 subjects (age 28.8 ± 2.3 years, range 25-35). These participants completed three to five 10-min long EEG-fMRI runs (mean 4.9 ± 0.6), including on average 29.3 ± 10.3 min of NREM sleep (range 13.5-53.0 min; N1 = 58.6 ± 25.2%; N2 = 41.2 ± 24.9%; N3 = 0.2 ± 0.8%; Figure S1). A total of 2021 slow waves (118.9 ± 79.4 per participant) were automatically detected using validated algorithms (Mensen et al., 2016; Siclari et al., 2014) based on the identification of consecutive signal zero-crossings (0.25-1.0 s half-wave duration, no amplitude thresholds; see Material and Methods) and were included in subsequent analyses.

### Changes in cortical and subcortical hemodynamic activity

A voxel-wise regression analysis (Figure 2) was used to identify significant changes in brain activity associated with the occurrence of NREM slow waves. We found that slow waves were associated with two distinct blood-oxygen-level-dependent (BOLD) responses (q < 0.01, FDR corrected with an additional minimum cluster threshold of 50 voxels). Specifically, significant signal increases were found in subcortical structures comprising the bilateral thalamus, the cerebellum, mainly posterior portions of the brainstem, and the right caudate nucleus (Figure 3A and Figure S2A, Table 1). Significant hemodynamic decreases were instead observed at cortical level in the bilateral somatomotor cortex, visual cortex and posterior insula, as well as in the left parahippocampal gyrus, in the right hippocampus and in the left parieto-occipital sulcus (Figure 3B and Figure S2B, Table 1).

**Figure 1.**
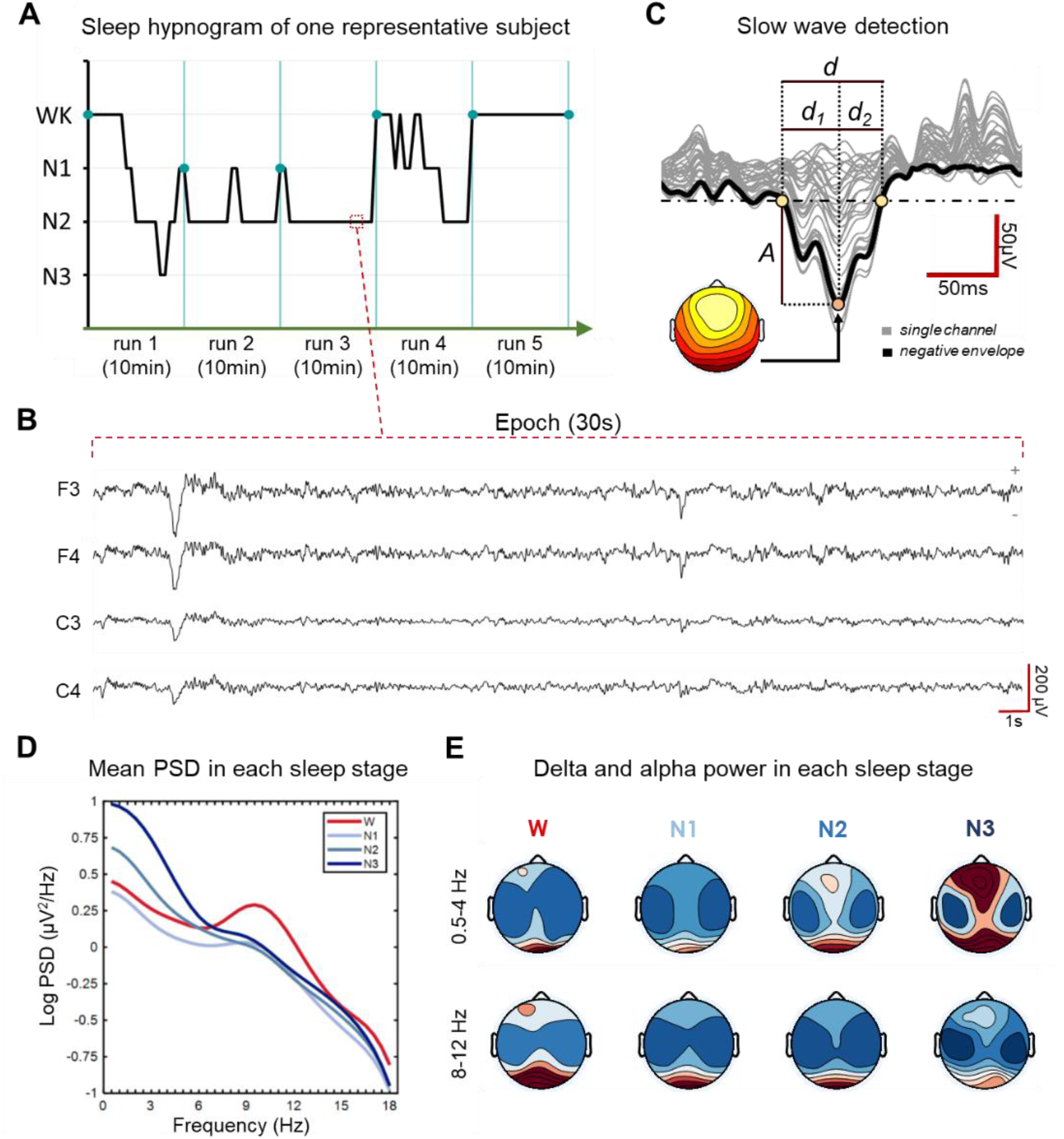
Scoring and analysis of EEG recordings. (A) Sleep hypnogram of one study participant. Vertical lines mark the end ofEEG-fMRI acquisition runs. (B) EEG traces of a N2 epoch of the same participant shown in panel A. (C) Schematic representation of the approach used to detect and characterize sleep slow waves. Negative half-waves were automatically detected through the identification of negative peaks comprised between consecutive signal zero-crossings. A = amplitude; d = half-wave duration; d_1_ = time from first zero-cross to negative peak; d_2_= time from negative peak to second zero-cross. (D) Mean power spectral densities (PSD) obtained across all studied volunteers for wakefulness (W), N1, N2, and N3 sleep. PSD values were computed in two frontal (F3, F4) and two central (C3, C4) electrodes and then averaged. (E) Changes in delta and alpha power across sleep stages were evaluated to verify the accuracy of sleep scoring procedures. As expected, all sleep stages are characterized by a reduction in alpha activity (8-12 Hz) with respect to wakefulness, while delta activity (0.5-4 Hz) increase from N1 to N3 sleep.

**Figure 2.**
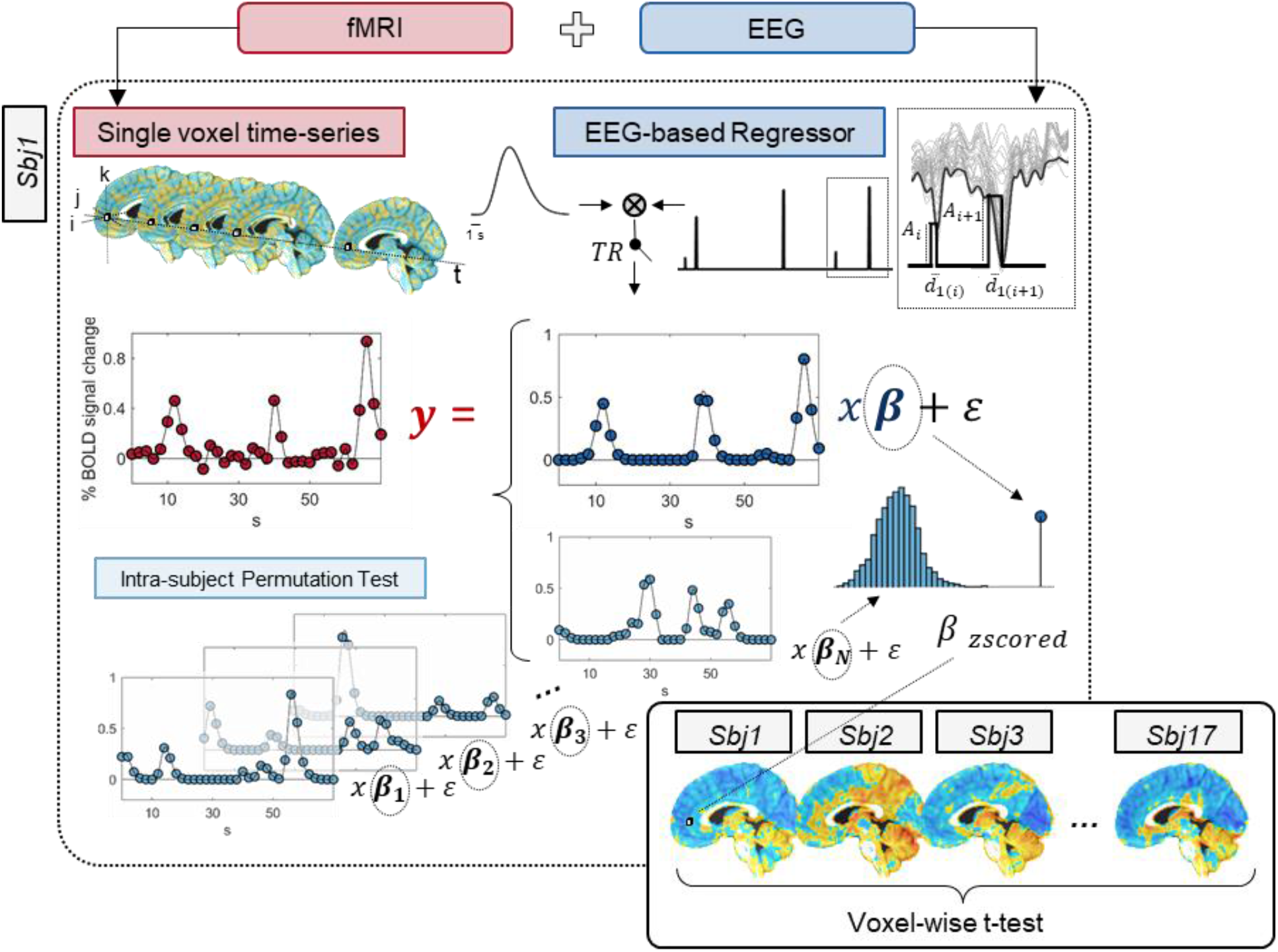
Schematic representation of procedures applied for the voxel-wise regression analysis. Each slow wave was modelled as a square wave with onset-time corresponding to the timing of the first zero-crossing, height equal to the absolute value of the maximum negative amplitude of the slow wave (A), and duration corresponding to the duration of the descending phase of the wave (d_1_). The obtained regressor was then convoluted with a standard gamma hemodynamic response function (HRF), down-sampled to the fMRI sampling rate and used to regress the BOLD time-series from each voxel. At single-subject level, beta-values of each voxel were converted into z-scores calculated with respect to a null distribution obtained by re-computing the regression after shuffling the timing of individual slow waves (nPerm =1000). A one-sample t-test was performed to assess statistical significance at group-level.

**Figure 3.**
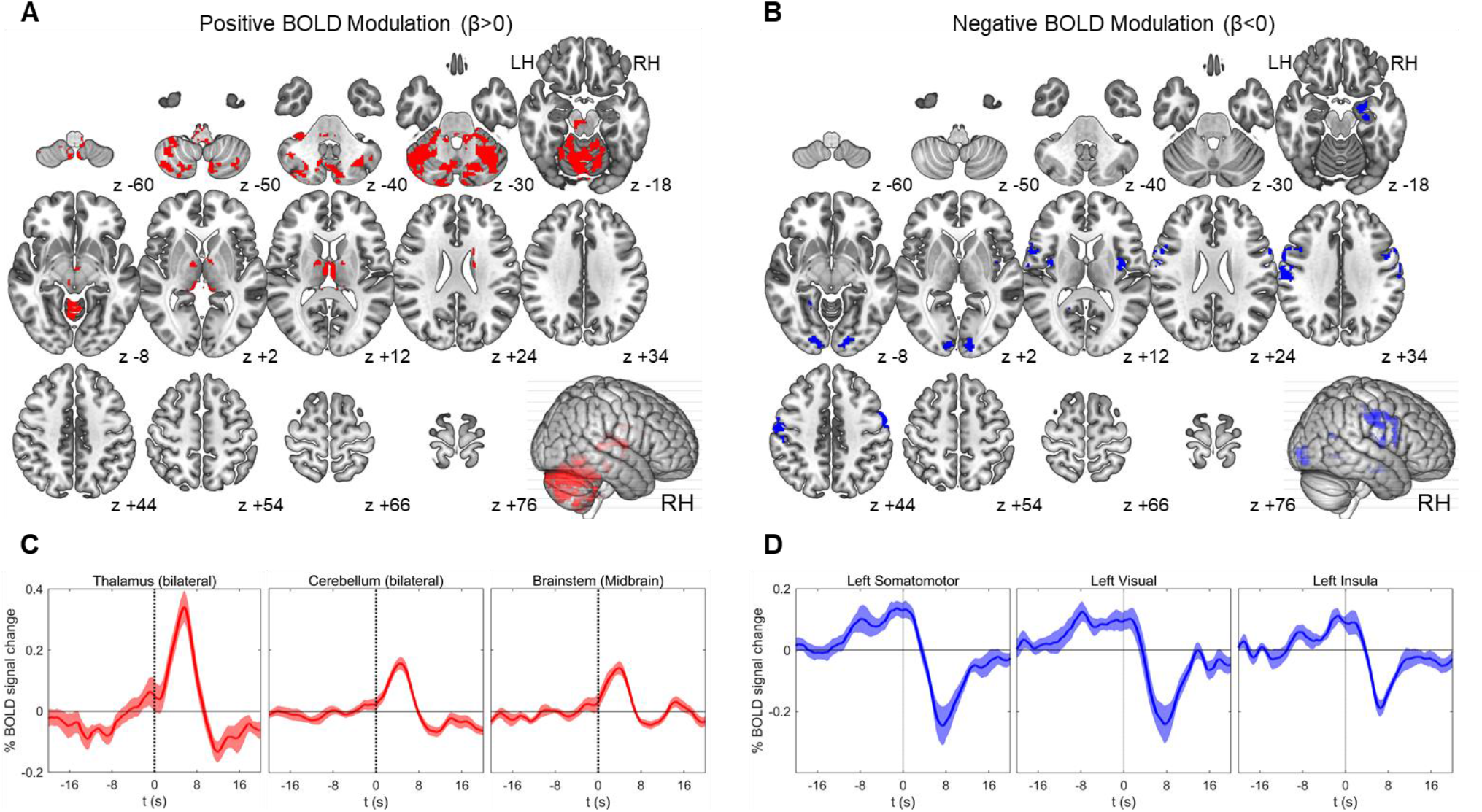
Results of the regression analysis. Brain structures associated with a significant (q < 0.01, cluster size ≥ 50 voxels) BOLD-signal (A) increase (red) or (B) decrease (blue). Brain images were generated using MRIcron (https://www.nitrc.org/projects/mricron). (C-D) Mean BOLD-signals (up-sampled to the EEG sampling rate) and relative standard errors for three subcortical (C) and three cortical (D) of the identified significant clusters. Time t = 0 s corresponds to slow-wave onset.

**Table 1.** Results of regression analysis. Brain areas showing a significant BOLD-signal increase (positive) or decrease (negative) in relation to the occurrence of sleep slow waves (q < 0.01, cluster size ≥ 50 voxels). The table includes the number of voxels and the coordinates of the center of mass of each significant cluster in standard MNI space.

| Signal change | Region | Voxels | CM x | CM y | CM z |
| --- | --- | --- | --- | --- | --- |
| Positive | Brainstem (Medulla) | 120 | -0.8 | -38.5 | -46.7 |
|  | Brainstem (Midbrain) | 95 | -7.3 | -25.3 | -15.7 |
|  | Caudate Nucleus | 76 | +17.9 | -0.8 | +25.5 |
|  | Cerebellum | 6322 | -5.2 | -62.7 | -29.2 |
|  | Cerebellar Tonsil | 121 | -2.4 | -54.7 | -61.3 |
|  | Medial Thalamus | 734 | +1.0 | -19.0 | +7.6 |
| Negative | Left Posterior Insula | 64 | -39.0 | -4.8 | +10.2 |
|  | Right Posterior Insula | 123 | +37.9 | -5.3 | +12.1 |
|  | Right Hippocampus | 135 | +24.8 | -10.1 | -18.9 |
|  | Left Parahippocampus | 50 | -22.0 | -54.6 | -4.9 |
|  | Left Parieto-Occipital Sulcus | 64 | -17.0 | -58.6 | +17.5 |
|  | Left Somatomotor Cortex | 800 | -57.4 | -7.4 | +32.5 |
|  | Right Somatomotor Cortex | 279 | +58.4 | -2.1 | +34.5 |
|  | Left Visual Cortex | 176 | -15.5 | -90.5 | -5.7 |
|  | Right Visual Cortex | 268 | +13.5 | -90.8 | -3.5 |

An evaluation of the average temporal profile of the hemodynamic response in each significant cluster revealed that the onset of subcortical BOLD-signal increases was temporally aligned with the onset of the sleep slow wave (t ≈ 0 s). These initial increases were followed by a slow negative BOLD-signal deflection that was especially evident in the thalamus and cerebellum. On the other hand, the strong negative signal deflection in neocortical areas was delayed by 2-4 s with respect to slow-wave onset, and was preceded by a slow, positive BOLD-signal deflection that started 4-12 s prior to slow-wave onset.

### Thalamic involvement in sleep slow waves

Previous work in animal models suggested that especially centro-medial thalamic nuclei may have a key role in the modulation of sleep slow waves (Gent et al., 2018b). In order to investigate whether this may be true also in humans, here we analyzed and characterized the anatomical and functional nature of thalamic portions recruited during the occurrence of human sleep slow waves (Figure 4). First, we determined the percentage of activated voxels that fell within specific thalamic substructures identified using a probabilistic atlas of the human thalamus based on diffusion-weighted imaging (Najdenovska et al., 2018). For both the left and the right thalamus, we found that significant BOLD-signal changes especially involved medial nuclei located anteriorly, centrally and posteriorly (Figures 4D-E). In particular, activated portions of the thalamus included ~40% of anterior thalamic nuclei (A), ~30% of posterior nuclei (medial pulvinar, PuM; central lateral nucleus, CL; lateral posterior nucleus, LP), ~20% of ventral anterior nuclei (VA) and ~18% of mediodorsal nuclei (MD).

**Figure 4.**
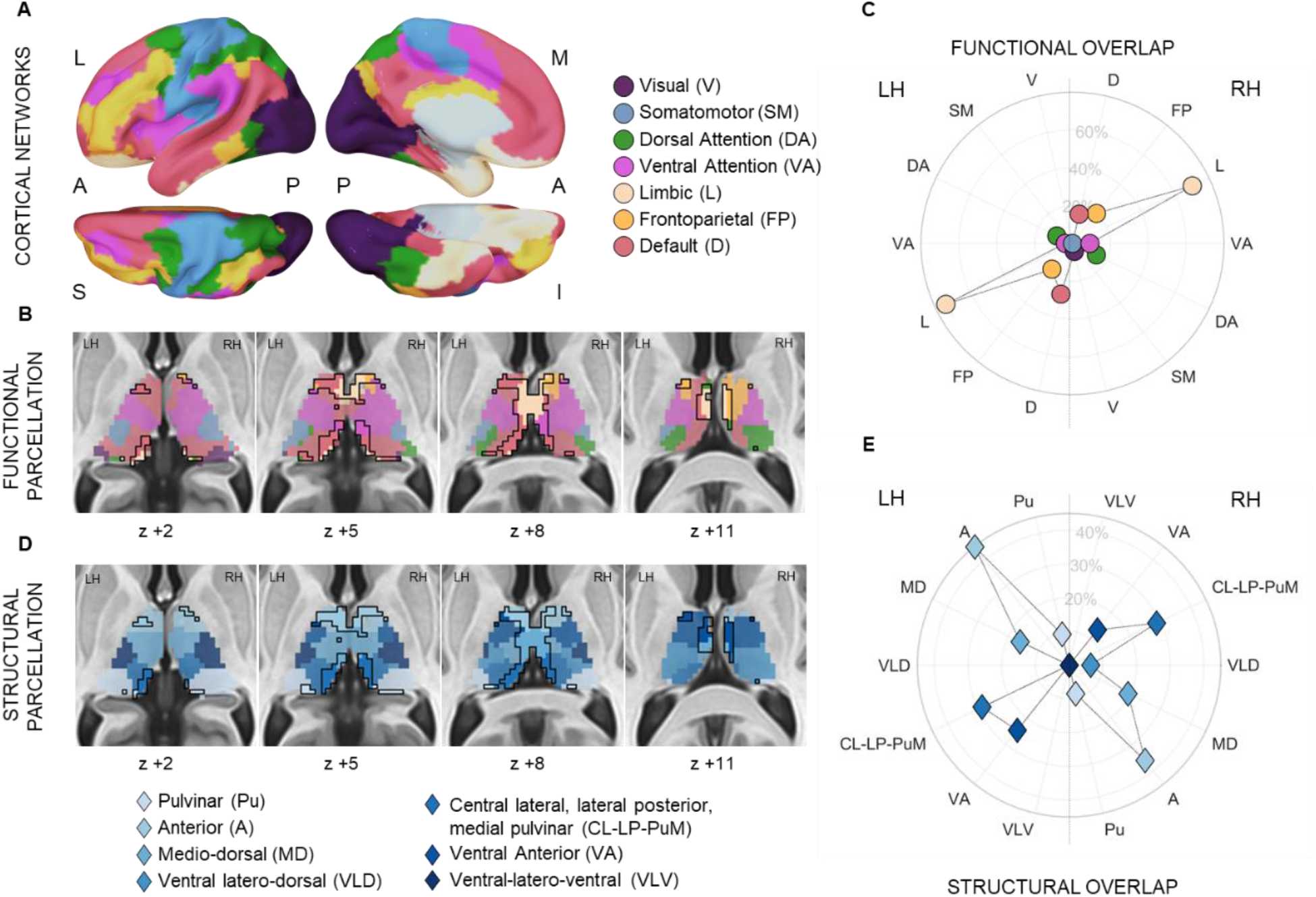
Thalamic involvement in sleep slow waves. (A) The seven canonical networks used to determine the preferential connectivity of each thalamic voxel. (B) Parcellation of the thalamus based on the preferential functional connectivity of each voxel. (C) Radial plot showing the proportions of activated voxels having a preferential connectivity with each of the seven canonical networks with respect to the totality of thalamic voxels. (D) Parcellation of the thalamus based on the Najdenovska atlas. (E) Radial plot showing the proportions of activated voxels attributed to each anatomical area with respect to the totality of thalamic voxels. In (B) and (D), the portion of the thalamus activated in association with the occurrence of sleep slow waves (q < 0.01, cluster size ≥ 50 voxels) is highlighted and enclosed with a black line.

Second, we evaluated the functional organization of activated thalamic portions by determining the preferential functional connectivity of each voxel with each of the seven main canonical brain networks (Hwang et al., 2017; Yeo et al., 2011): visual, somatomotor, dorsal attention, ventral attention, limbic, fronto-parietal and default mode (Figure 4A-B). We found that the portion of thalamus recruited during sleep slow waves was especially connected with areas of the limbic system (Figure 4C). Indeed, the activated thalamic cluster included ~70% of all thalamic voxels showing a preferential connectivity with this particular network. Other represented networks included the default mode network (~22%) and the frontoparietal network (~20%).

### Cerebellar involvement in sleep slow waves

Changes in cerebellar activity have been previously reported as a function of sleep stage as well as in association with the occurrence of sleep slow waves (for a detailed review see Canto et al., 2017). Yet, the determinants of sleep-dependent changes in cerebellar activity and the possible role of the cerebellum in sleep regulation are still largely unknown. In order to advance current knowledge on these key aspects, here we performed a detailed evaluation of cerebellar activity related to the occurrence of NREM slow waves (Figure 5). As described above for the thalamus, we first determined the percentage of activated voxels that fell within specific cerebellar areas identified using the SUIT anatomical atlas (Diedrichsen et al., 2009, 2011). The results of this analysis are presented in Figure 5C-D and indicate a broad involvement of the vermis and of the superior portions of both cerebellar hemispheres. Then, we evaluated the functional relationship between activated cerebellar portions and the seven canonical brain networks (Figure 5A-B). We found that the fractions of cerebellum involved in sleep slow waves were especially connected with areas of the somatomotor network. In fact, the activated cluster included ~45% of all cerebellar voxels showing a preferential connectivity with this particular network. Other represented networks included the frontoparietal network (~34%) and the ventral attention network (~28%).

**Figure 5.**
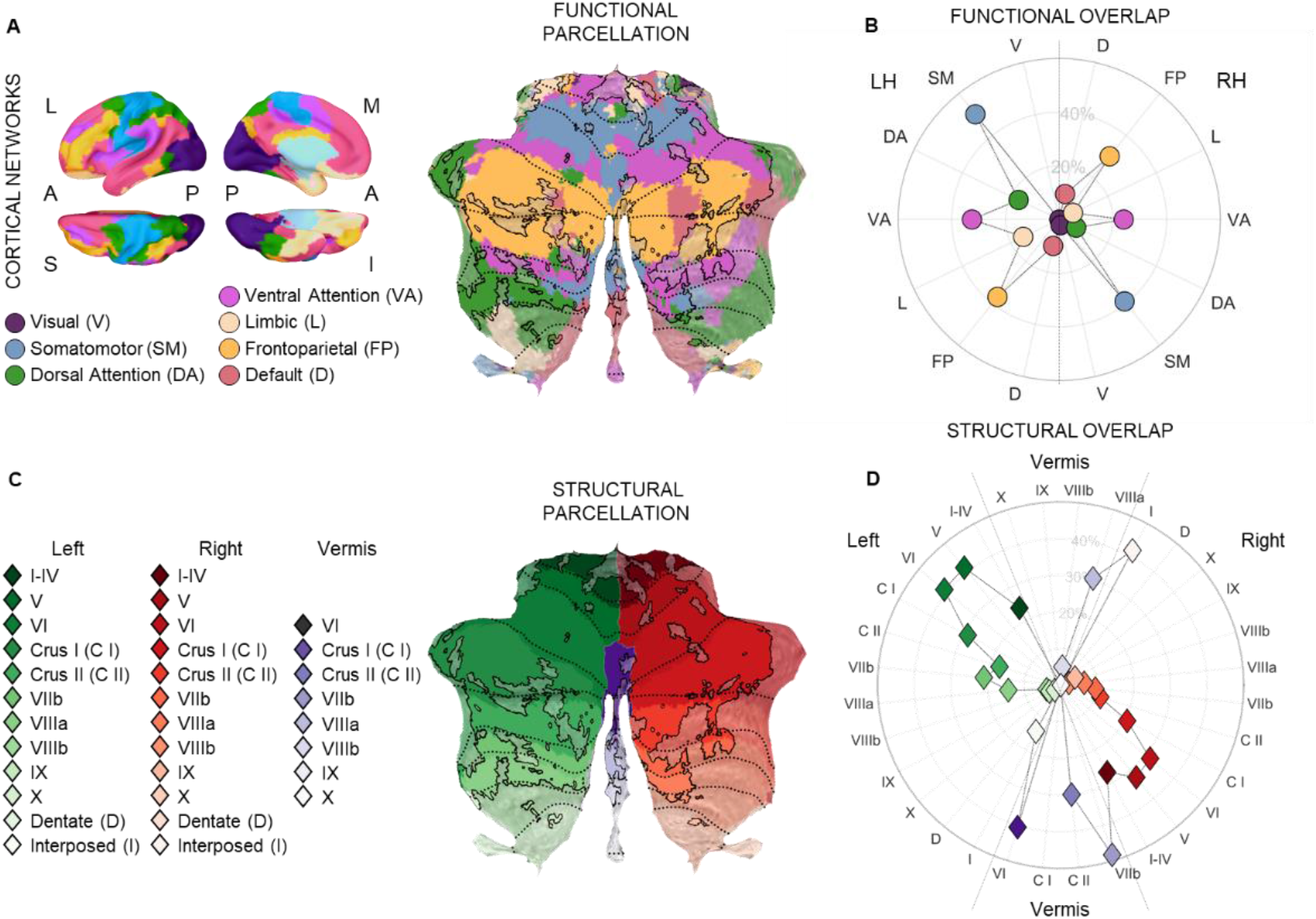
Cerebellar involvement in sleep slow waves. (A) The seven canonical networks and the correspondent parcellation of the cerebellum based on the preferential connectivity of each voxel. (B) Radial plot showing the proportions of activated voxels having a preferential connectivity with each of the seven canonical networks with respect to the totality of cerebellar voxels. C) Anatomical parcellation of the cerebellum based on the SUIT anatomical atlas. (D) Radial plot showing the proportions of activated voxels attributed to each anatomical area with respect to the totality of cerebellar voxels. In (A) and (C), the portion of the cerebellum activated in association with the occurrence of sleep slow waves (q < 0.01’ cluster size ≥ 50 voxels) is highlighted and enclosed with a black line.

### Cortical involvement in sleep slow waves

Macro-scale hd-EEG studies in humans showed that slow waves are not stationary events, but instead propagate at cortical level through anatomically connected pathways (Avvenuti et al., 2020; Massimini et al., 2004; Murphy et al., 2009). This propagation of electrophysiological slow waves may determine a relative variability in the timing of hemodynamic changes across cortical regions. Of note, BOLD-changes occurring ahead or delayed with respect to the actual onset of slow waves, could be missed by conventional fMRI analyses (Mitra et al., 2015). In line with this possibility, a preliminary evaluation of the mean BOLD-signal time-courses within areas of the seven canonical networks showed similar patterns of changes in all regions, thus suggesting that most cortical areas may actually present some level of modulation in association with the occurrence of sleep slow waves (Figure 6).

**Figure 6.**
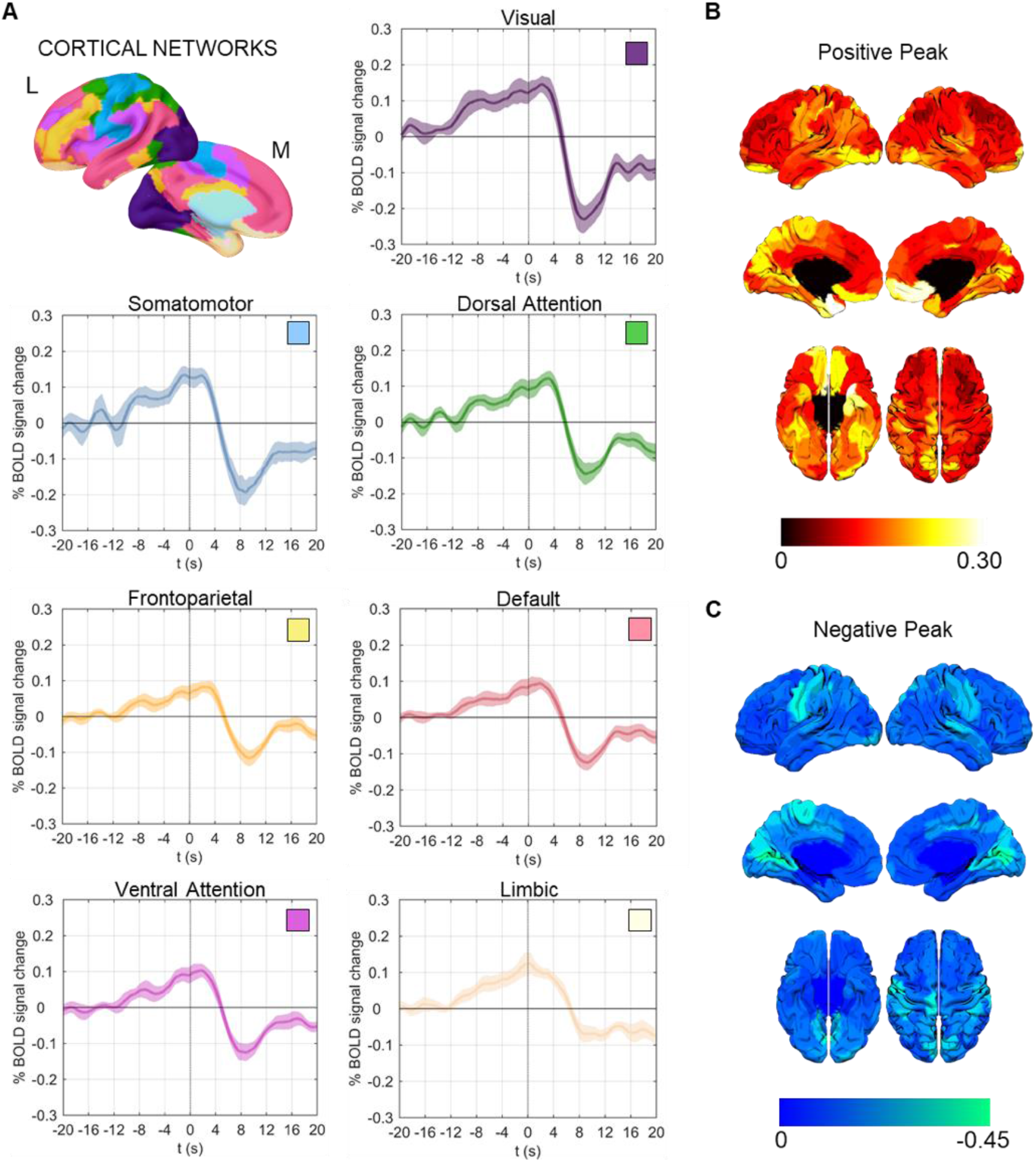
Slow-wave-related hemodynamic changes across the cortical mantle. (A) Mean BOLD-signal time-courses (± standard error) associated with sleep slow waves in the seven canonical networks. Time t = 0 s corresponds to slow-wave onset (positive-to-negative zero crossing). (B) Amplitude of the positive (top) and negative (bottom) peaks in each of the 200 ROIs of the Schaefer atlas. Brain surface plots were generated using the Surf Ice software (https://www.nitrc.org/projects/surfice/). The largest negative amplitudes were found in primary cortices, while the largest positive amplitudes were also found in areas of the limbic network.

In light of these observations, further analyses were performed to investigate whether hemodynamic cortical changes showed a relative propagation similar to the one described for electrophysiological slow waves. Specifically, we first computed the average BOLD-signal across slow waves and subjects within 200 cortical regions of interest (ROIs) of the Schaefer functional atlas (Schaefer et al., 2018; Figure 6B shows the relative amplitude of positive and negative BOLD-signal peaks for all ROIs). Then, a cross-correlation analysis was performed between a seed time-series corresponding to the left somatomotor cortex and the time-series of all brain ROIs. This investigation confirmed that most brain areas showed similar slow-wave-dependent hemodynamic changes, which however occurred at different delays in distinct brain regions (Figure 7; also see Figure S3). In particular, the lowest delays were found in areas encompassing the somatomotor cortex, the premotor-prefrontal cortex and the anterior insula, while the highest delays were found in the inferior and lateral occipital and temporal cortex. These observations were substantially confirmed through a network-level analysis based on seven large bilateral ROIs. In fact, the lowest latency was found in the somatomotor network, followed by the ventral attention (+ 0.07 s) and the dorsal attention (+ 0.33 s) networks. The highest delays were instead found in the visual (+ 0.80 s) and limbic (+ 1.00 s) networks.

**Figure 7.**
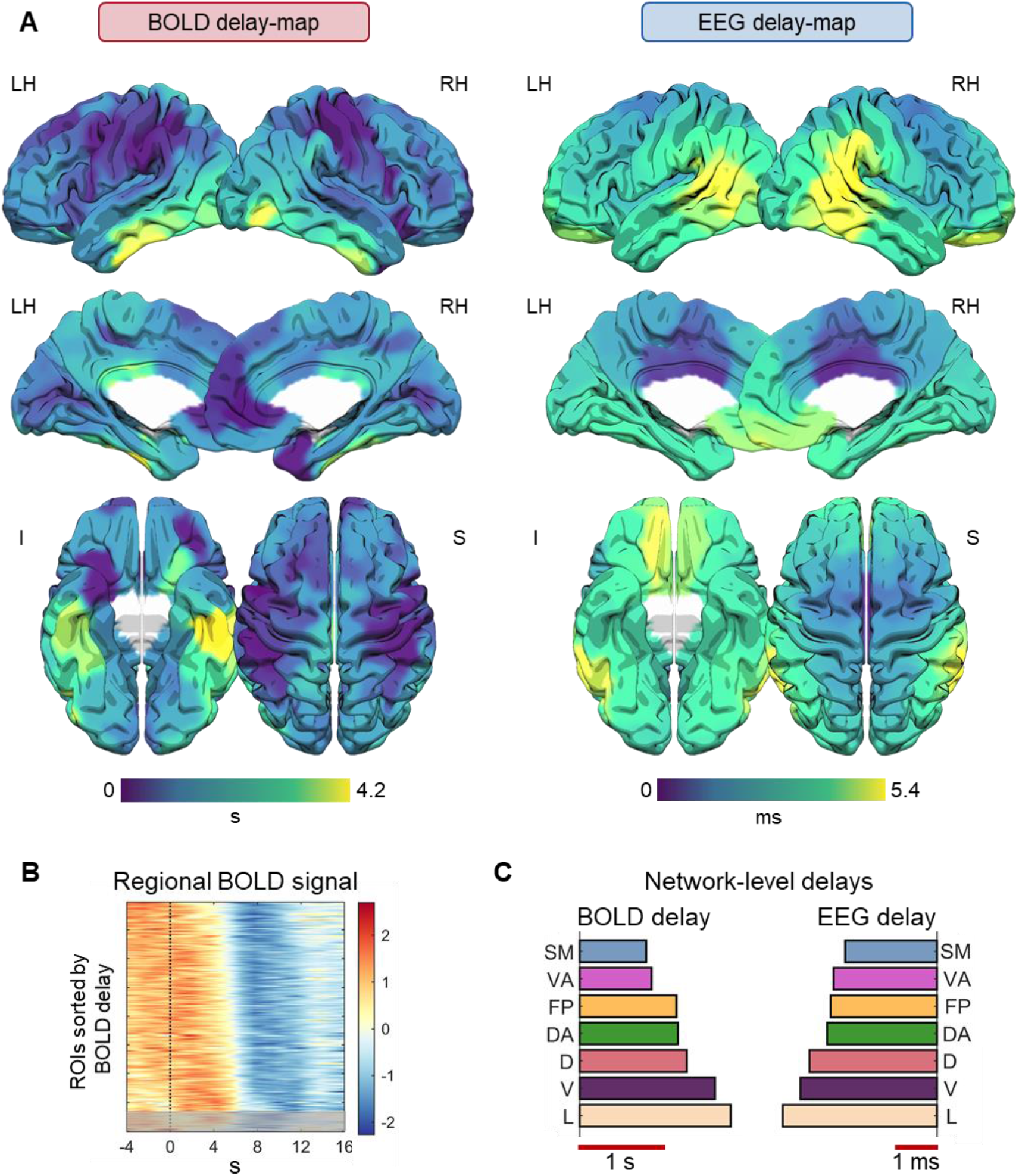
Hemodynamic and electrophysiological signal propagation during sleep slow waves. (A) Mean cortical delay-maps. Brain surface plots were generated using the Surf Ice software (https://www.nitrc.org/projects/surfice/). Here the yellow color indicates a high delay, while the dark-blue color indicates a low delay. All the 200 ROIs of the Schaefer atlas are shown. It is important to note that these maps reflect a mean propagation pattern, which may not correspond to the propagation pattern of each individual slow wave. (B) Mean hemodynamic brain activity changes (z-score) in the 200 ROIs, sorted according to relative delay values. ROIs characterized by a non-significant cross-correlation with the seed region were put in the bottom section of the image and are covered by a gray shadow. (C) Mean propagation delays computed for the seven (bilateral) canonical networks for BOLD (left) and EEG (right) signals. SM: somatomotor, VA: ventral attention, FP: fronto-parietal, DA: dorsal attention, D: default mode, V: visual, L: limbic.

Interestingly, while spanning a much broader time-frame (in the order of seconds vs. tens to few hundred of milliseconds), the fMRI delay-map was strikingly reminiscent of the one obtained from the analysis of slow-wave propagation in source-level hd-EEG data (Murphy et al., 2009). In order to allow a direct comparison between the hemodynamic and the electrophysiological delay-maps, here we re-analyzed NREM-sleep hd-EEG recordings (256 electrodes, EGI-Philips; 500 Hz sampling frequency) obtained in a distinct set of 12 healthy adult volunteers (see Materials and Methods). In particular, slow waves were automatically detected in the first 30 min of N2/N3 sleep, and their regional propagation delays were computed in source space using a previously described approach (Murphy et al., 2009; Siclari et al., 2014). In line with results obtained for hemodynamic changes, electrophysiological slow waves showed the lowest latency in areas encompassing somatomotor cortex, insula and premotor areas, while the highest delay was observed in temporal and occipital regions. A correlation analysis at ROI-level (N=182) confirmed the existence of a significant association between hemodynamic and electrophysiological delay-maps (Pearson’s ρ = 0.264; p = 0.0003; Figure S4). The same analysis performed at network-level (N=7) also yielded a strong correlation with ρ = 0.893 (p = 0.007; Figure 7C), suggesting that the relatively modest correlation coefficient obtained at ROI-level could be in part ascribed to inaccuracies in source estimation.

### Relationship between slow-wave amplitude and BOLD-signal variations

Finally, we investigated whether variations in the properties of electrophysiological slow waves corresponded to variations in the profiles of regional hemodynamic responses. A correlation analysis between slow wave amplitude and point-by-point BOLD-signal changes revealed a significant positive relationship between EEG amplitude and the absolute magnitude of the observed positive subcortical and the negative cortical signal-deflections (q < 0.01; Figure 8 and Figure S5). Of note, the small positive BOLD-signal deflection observed in cortical areas immediately before or around slow-wave onset, and the late negative component observed in subcortical structures were also significantly modulated by slow wave amplitude. Interestingly, significant positive correlations were found up to 4 s before slow-wave onset in thalamus, somatomotor cortex, insula and hippocampus, in line with previous evidence indicating a possible central role of these regions in slow wave generation (Murphy et al., 2009). The evaluation of hemodynamic changes for distinct slow-wave amplitude percentile classes (*A_1_*: 0-33; *A_2_*: 33-66; *A_3_*: 66-100) confirmed that larger slow waves were associated with larger BOLD-signal changes, and also revealed a relative modulation of the latency of such changes (trend analysis, p < 0.01). In fact, larger slow waves were associated with larger hemodynamic variations that occurred earlier in time, while smaller slow waves were associated with smaller hemodynamic changes that tended to be shallower and to peak later in time (Figure 8 and Figure S6).

**Figure 8.**
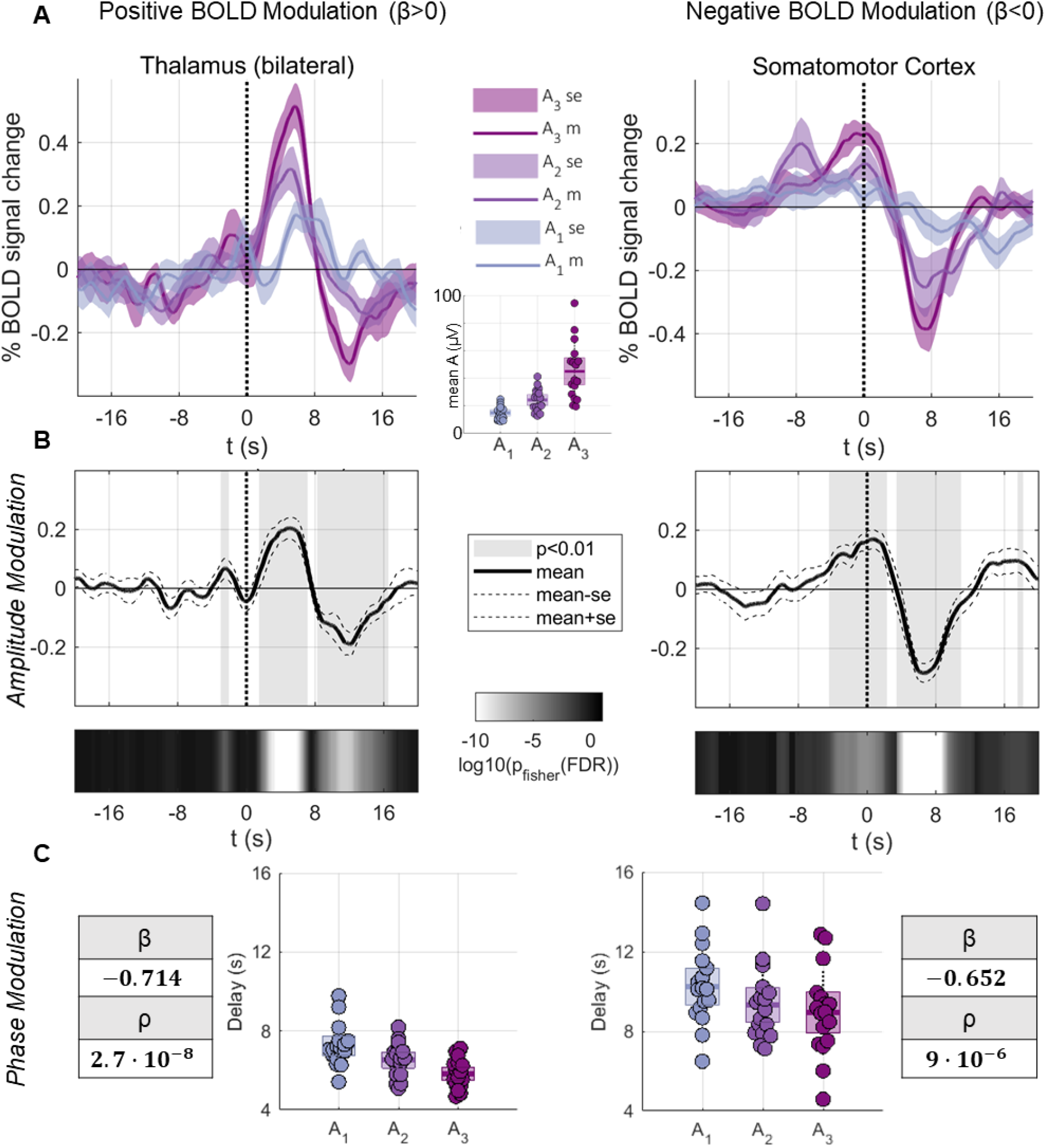
Relationship between EEG slow-wave properties and regional hemodynamic changes. (A) Temporal evolution of BOLD-signals (averaged across subjects), for three classes of slow waves defined based on amplitude percentiles (*A*_1_: 0-33; *A*_2_: 33-66; A_3_: 66-100). (B) Point-by-point correlation between slow-wave amplitude and BOLD-signal change for representative subcortical (thalamus) and cortical (visual cortex) structures. The gray shadowed areas indicate significant effects with q<0.01. (C) Relative delays of the positive (for thalamus) and the negative (for visual cortex) BOLD-signal peaks for each of the three wave classes. A regression was performed to determine whether a relationship existed between slow wave amplitude and the relative latency of hemodynamic peaks. Correspondent beta and p-values are reported in the gray boxes.

## Discussion

To the best of our knowledge this is the first study to provide an in-depth characterization of cortical and subcortical brain-activity patterns associated with sleep slow waves in humans. Our results can be summarized in four main findings. First, we showed that sleep slow waves are associated with significant BOLD-signal increases in subcortical structures, including the brainstem, thalamus and cerebellum, whereas in cortical areas a slow BOLD-signal increase precedes slow-wave onset and is followed by a prominent decrease. Second, we showed that positive subcortical changes occur within specific portions of thalamus and cerebellum, including medial thalamic nuclei having a strong connectivity with bilateral limbic functional networks, and cerebellar areas having a preferential connectivity with the somatomotor network. Third, we showed that cortical hemodynamic responses occur at different delays across a broad extent of the cortical mantle, with the shortest delay observed in somatomotor areas and the highest delay in temporal and lateral occipital areas. Importantly, the BOLD-signal propagation pattern substantially mirrors the spreading of electrophysiological slow waves. Finally, by investigating the relationship between EEG slow waves and hemodynamic activity, we found that slow-wave amplitude is directly correlated to the magnitude and the latency of the hemodynamic BOLD-signal changes in both cortical and subcortical areas. Overall, these findings indicate that human slow waves are associated with a complex chain of interactions among subcortical and cortical brain structures, in line with evidence obtained in animal models.

### Sleep slow waves are associated with hemodynamic changes in brainstem and thalamus

Our results indicate that human sleep slow waves are associated with hemodynamic changes in the brainstem and the thalamus, including an initial signal increase followed by a late, less pronounced negative deflection. Of note, the thalamic activation especially involves midline nuclei presenting a strong connectivity with the limbic network - the so-called ‘limbic thalamus’ (Colavito et al., 2015; Vertes et al., 2015). At the brainstem level, two main clusters of activation were found in the posterior midbrain and medulla. The posterior brainstem contains the *reticular formation* (Brown et al., 2012; Moruzzi and Magoun, 1949), which is known to send projections to the thalamic reticular nucleus and the midline and intralaminar thalamic nuclei (Mountcastle and Poggio, 1974; Steriade and Glenn, 1982). This may explain the co-activation of these structures in association with sleep slow waves.

The involvement of brainstem and medial thalamus in human slow waves is consistent with previous findings in animal models (Gent et al., 2018a; Neske, 2016). Indeed, while the *slow oscillation* underlying the generation of sleep slow waves is traditionally regarded as a prominently cortical phenomenon due to its persistence after thalamic lesion (Steriade et al., 1993) and in deafferented cortical slabs (Timofeev et al., 2000), accumulating evidence indicates a direct involvement of the brainstem and the thalamus in modulating important properties of NREM slow waves. In particular, these structures have been suggested to play a fundamental role in triggering and synchronizing the *up-state* at each cycle of the *slow oscillation* (Amzica and Steriade, 1995; Contreras and Steriade, 1995; Eschenko et al., 2012; Mena-segovia et al., 2008; Schweimer et al., 2011). Consistent with this view, previous work showed that cortical *up-states* are preceded by spontaneous thalamic spikes (Gent et al., 2018a; Sheroziya and Timofeev, 2014; Slézia et al., 2011; Ushimaru and Kawaguchi, 2015) and may be induced through electrical or optogenetic stimulation of the thalamus (David et al., 2013; Gent et al., 2018a; Honjoh et al., 2018; Poulet et al., 2012). Moreover, thalamic deafferentation of the cortex leads to a significant reduction in the frequency of *slow oscillations* (David et al., 2013; Lemieux et al., 2014). Our results support the possibility of a similar regulating role of subcortical structures during human slow waves. Moreover, the specific recruitment of limbic-related thalamic portions is congruous with evidence indicating a key-role of NREM slow waves in learning and memory consolidation (Diekelmann and Born, 2010; Miyamoto et al., 2017; Tononi and Cirelli, 2014).

### Cerebellar involvement in human sleep slow waves

A mainly positive hemodynamic modulation similar to the ones found in the brainstem and the thalamus was also observed in the cerebellum. In fact, an association between sleep slow waves and cerebellar activity has been previously described both in animal models (Roš et al., 2009; Steriade et al., 1971a, 1971b) and in humans (Dang-Vu et al., 2008). In addition, during N2 sleep, changes in cerebellar activity have been found to accompany the occurrence of K-complexes (Jahnke et al., 2012). Evidence obtained in ketamine-anesthetized rats suggests that the neocortex may entrain the cerebellum during the alternation of *down-* and *up-states* (Rowland et al., 2010), while the cerebellum may in turn have a relevant role in the fine-tuning of cortical slow waves (Canto et al., 2017). Nevertheless, the mechanism and functional meaning of cerebellar recruitment during cortical *slow oscillations* is still to be fully understood. One intriguing possibility is that changes in cerebellar activity play a role in the consolidation of motor memories, and possibly of other cognitive skills (Canto et al., 2017). In line with this view, here we found that portions of the cerebellum activated during sleep slow waves especially include those connected with the somatomotor network and, to a lesser extent, to the spatially close frontoparietal and ventral-attention networks.

### Human slow waves are coupled with cortically propagating hemodynamic waves

At the cortical level, we found that slow-wave onset is associated with a negative BOLD-signal deflection that could reflect the occurrence of locally synchronized *down-states* characterized by neuronal silence. In particular, we observed significant cortical clusters displaying large BOLD-signal decreases in bilateral insula, somatomotor cortex and visual cortex, as well as in the right hippocampus, left parahippocampus and left parieto-occipital sulcus. However, similar hemodynamic changes were also found to occur at different delays across distinct brain areas, covering a broad extent of the cortical mantle. In fact, we found that the insula, somatomotor and premotor regions were among the areas with the lowest relative delay, while the highest delays were found in the occipital and temporal cortex. This finding is consistent with the known cortical propagation of electrophysiological slow waves. Indeed, previous work showed that most EEG slow waves have a well-defined and circumscribed origin that more often involve brain areas located around the somatomotor cortex and the insula (Avvenuti et al., 2020; Massimini et al., 2004; Menicucci et al., 2009; Murphy et al., 2009), from which they propagate toward more anterior and posterior areas. In line with this, here we showed that the regional delays of hemodynamic changes significantly correlate with the propagation delays of electrophysiological slow waves, thus implying a direct coupling between the two phenomena. This finding is especially interesting in light of recent evidence linking EEG slow waves, hemodynamic changes and CSF movements (Fultz et al., 2019). Indeed, while CSF movement was not directly assessed in the present work, our results suggest that slow-wave propagation could generate a hemodynamic gradient that may ultimately favour CSF flow and the clearance of metabolic wastes. Interestingly, this mechanism could also contribute to explain the recent observation of a direct relationship between slow wave activity (1-4 Hz, delta power) and glymphatic CSF movement (Hablitz et al., 2019). Indeed, here we found that the amplitude of electrophysiological slow waves is positively related to the magnitude and inversely related to the delay of cortical (and subcortical) BOLD-signal changes. Thus, larger slow waves are associated with stronger and faster hemodynamic changes that may lead to a more efficient mobilization of the CSF. This hypothesis would be consistent with previous reports of a direct link between alterations of slow wave activity and cognitive decline related to the accumulation of β-amyloid in the medial prefrontal cortex of older individuals (De Gennaro et al., 2017; Mander et al., 2015).

Why slow waves show the lowest delay, and thus, a more common origin, in insula and somatomotor and premotor cortex, is unclear. These regions are characterized by a strong noradrenergic innervation (Gaspar et al., 1989; Javoy-Agid et al., 1989; Lewis and Morrison, 1989) and may thus be particularly affected by the overall reduction of activating neuromodulators from wakefulness to NREM sleep. Moreover, the same areas also seem to represent a preferential origin for large and widespread slow waves (including K-complexes), whose generation/synchronization mechanism has been suggested to depend on phasic activity of ascending activating systems (Bernardi et al., 2018; Siclari et al., 2014). Another intriguing possibility is that the spatial distribution of slow-wave origin reflects a role of slow waves in the maintenance of sensory and motor disconnection during sleep (Funk et al., 2016). Possibly consistent with this interpretation, we found that the cortical negative signal deflection is preceded by a slow rise of the BOLD-signal, which peaks around the timing of slow wave onset. This BOLD-signal increase may reflect spontaneous or stimulus-induced increases in cortical activity that eventually trigger a response-suppression represented by the sleep slow wave. In fact, a similar mechanism has been suggested to subserve the sleep-protective function of stimulus-induced K-complexes (Andrillon and Kouider, 2020; Laurino et al., 2019, 2014; Riedner et al., 2011).

### Limitations

Present analyses were performed on EEG-fMRI data collected during an afternoon nap and most participants failed to reach deep, N3 sleep. It should be noted, however, that the relative sparsity of slow waves occurring in light sleep can be expected to minimize potential confounds related to the overlap of hemodynamic responses associated with slow waves occurring in close temporal sequence. This is especially important given the slow temporal evolution of the hemodynamic response and its relative propagation at cortical level. Given these considerations, future studies should verify whether trains of large slow waves that occur in N3 may involve partially distinct sets of brain regions or be associated with different response patterns. In particular, there is evidence for the existence of different subtypes of slow waves, including so-called K-complexes (type I slow waves) and delta waves (type II slow waves), which co-exist during NREM sleep and likely rely on different synchronization mechanisms (Bernardi et al., 2018; Siclari et al., 2014). It is therefore unlikely that all slow waves engage the whole set of brain regions in an identical manner. Indeed, not only each individual slow wave may recruit partially different areas depending on its specific origin and propagation pattern, but the same areas could be also engaged differently by distinct slow-wave subtypes.

### Conclusions

Overall, present results indicate that human slow waves are associated with complex patterns of hemodynamic changes, including both increases and decreases, and involving cortical as well as subcortical structures. These patterns of brain activity are consistent with theoretical accounts of the functions of sleep slow waves. In particular, the strong connectivity between activated thalamic nuclei and limbic structures is consistent with a role of sleep slow waves in memory processing (Miyamoto et al., 2017). Moreover, our results demonstrate coupled electrophysiological and hemodynamic fluctuations that orderly propagate from a preferential origin in centro-frontal cortical brain areas to more anterior and posterior regions. We hypothesize that these coupled propagation dynamics could have a direct role in the generation of gradients of CSF flows and in the clearance of metabolic wastes (Fultz et al., 2019). The mechanism and function of the slow-wave-dependent electrophysiological-hemodynamic coupling undoubtedly deserves future investigations due to its possible implication in pathological conditions.

## Materials and Methods

### Participants

Twenty healthy adult volunteers (age 29.7 ± 3.9 years, range 25-42; 11 females; all righthanded) participated in the study. A clinical interview was performed to rule out history or presence of any disorder that could significantly affect brain function. Then, recruited participants underwent simultaneous EEG-fMRI recording during an afternoon nap opportunity. In order to facilitate transition into sleep during the scan session, subjects were asked to wake-up 1 to 2 hours earlier than usual in the morning of the experiment and to restrain from consuming caffeine-containing beverages in the few hours that preceded the scan session. On average, participants reported to have slept for 4.9 ± 1.5 h the night before the experiment, which corresponded to 66.2 ± 26.3 % of their usual sleep time.

As detailed below, we also re-analyzed resting state (rs-)fMRI and hd-EEG data collected in two independent samples of healthy adult individuals. Specifically, rs-fMRI data was obtained in a group of 28 adult subjects (age 25.0 ± 5.3 years, range 19-43; 14 females; 14 left-handed). Data acquisition was performed using the same MRI scanner and acquisition parameters comparable to those employed in the present study (see below). The hd-EEG data consisted of overnight sleep recordings obtained at the University Hospital of Lausanne (Switzerland) in a distinct set of 12 healthy adult volunteers (age 25.5 ± 3.7 years, 6 females) who participated in a larger project aimed at exploring the effect of visual experience on sleep slow waves (Bernardi et al., 2019a). Here we only included data from the control condition, in which subjects remained in the sleep laboratory and watched movies of their choice (selected from a pre-defined list) from 3 to 8PM.

All experiments were conducted under protocols approved by the respective Local Ethical Committees, in accordance with the ethical standards of the 2013 Declaration of Helsinki. Written informed consent was obtained from all participants.

### Data acquisition

All study participants underwent simultaneous EEG, ECG and fMRI recordings during an afternoon nap opportunity (2:30-4:00PM). Custom-made foam pads were used to improve the comfort of the subjects inside the coil and minimize possible head movements. The participants received instruction to relax and try to sleep in the scanner. The study was interrupted after acquisition of five 10-min fMRI runs or when the participant started to feel uncomfortable in the scanner and/or felt unable to fall asleep.

The EEG was recorded using an MR-compatible EEG cap (Micromed, Mogliano Veneto, Italy) including 32 electrodes online-referenced to FCz (1,024 Hz sampling frequency). The electrode-skin impedance was brought below 10 KΩ at the beginning of the scan session. During simultaneous EEG-fMRI acquisition, electrophysiological data were transmitted through a fibreoptic cable from the high-input impedance amplifier (22 bits resolution, with range ± 25.6 mV) in the scanner room to a computer located outside. Before transmission, the signal was band-pass filtered between 0.15 and 269.5 Hz by an anti-aliasing hardware band-pass filter.

Functional and anatomical data were acquired using a 3T Philips Achieva MR-scanner. T2*-weighted gradient-echo echoplanar sequences were used to acquire functional data from 39 axial contiguous slices (300 volumes per run; TR = 2000 ms; TE = 35ms; FA = 80°; voxel size: 3×3×3 mm). A high-resolution T1-weighted MPRAGE anatomical image was also obtained at the end of the experimental session. The volume consisted of 170 sagittal slices (TR = 9.9 ms; TE = 4.6 ms; in plane matrix = 256×256; voxel size = 1×1×1 mm).

### EEG data preprocessing and analysis

All EEG recordings were preprocessed in MATLAB (The MathWorks, Inc.) using EEGLAB (Delorme and Makeig, 2004). The FMRIB-plugin (Iannetti et al., 2005; Niazy et al., 2005) was used to remove fMRI-related artifacts from EEG data through the following steps: removal of MR gradient artifacts based on average artifact subtraction (AAS), subtraction of the principal components (PC) of artifact residuals (Optimal Basis Sets method; OBS) and adaptive noise cancelation (ANC); signal down-sampling to 256 Hz; automated detection of QRS complexes in ECG channel followed by removal of ballistocardiographic artifacts (i.e., artifacts caused by cardiac pulse-related movement of the scalp electrodes inside the magnetic field; Allen et al., 2000) based on subtraction of PCs of artifact residuals (OBS). Of note, all automated QRS detections were visually inspected and wrong or missing markers were manually corrected using custom-made MATLAB functions. Finally, EEG recordings were band-pass filtered between 0.5 and 18 Hz and an Independent Component Analysis (ICA) was applied to remove any residual activity of artifactual origin, including artifacts related to eye movements and muscular activity. Bad channels were visually identified, rejected and interpolated using spherical splines from the activity of the nearest sensors.

The EEG signal was re-referenced to the average of channels T5 and T6, and sleep scoring was performed over 30-s epochs according to standard criteria (Iber et al., 2007). Epochs containing residual artifactual activity were manually marked through visual inspection and excluded from further analyses as detailed below. For the automated detection of sleep slow waves a negative-going signal envelope was calculated by computing the average of the fourth, third and second most negative samples across all electrodes (Mensen et al., 2016). Of note, the most negative sample was discarded to minimize the potential impact of any residual large-amplitude artifactual activity in isolated electrodes. The resulting signal was baseline corrected (zero mean-centered) prior to the application of a negative-half-wave detection procedure based on the identification of consecutive signal zero-crossings (Riedner et al., 2007; Siclari et al., 2014). Differently from commonly applied channel-by-channel detection approaches, this method allows to identify both local and widespread slow waves and to define a unique time reference (across electrodes) for each negative wave (Mensen et al., 2016). Only negative half-waves detected during NREM sleep epochs (N1/N2/N3) and with a duration comprised between 0.25 s and 1.0 s (full-wave period 0.5–2.0 s, corresponding to a 0.5-2.0 Hz frequency range) were selected for subsequent analyses. An amplitude threshold was not applied in order to allow investigating the relationship between slow wave amplitude and BOLD-signal changes. The timing of the first zero-cross (from positive to negative) was used as a reference to mark the beginning of each slow wave (slow-wave onset). Moreover, the amplitude (*A*; unit: μV), defined as the absolute value of the maximum negative peak, and the zero-cross-to-negative-peak time (*d*_1_; unit: s) of all detected half-waves were computed and stored for further analyses.

### MRI data preprocessing and analysis

Functional MRI data were preprocessed using *AFNI* (Cox, 2012, 1996). First, signal outliers were removed from single-voxel time-series *(3dDespike)* and the time-shift related to slice acquisition was corrected *(3dTshift).* Data from different runs were then registered to a reference volume for motion correction *(3dvolreg)* and spatially smoothed *(3dBlurToFWHM)* with a 6 mm full-width-at-half-maximum (FWHM) Gaussian kernel. For each fMRI run, the signal of each voxel was converted to percent BOLD-signal change with respect to the mean BOLD-signal across the corresponding run. Additional preprocessing steps were performed to remove potential artifactual components in the fMRI signal. In particular, the BOLD-signal from each voxel was cleaned by regressing-out *(3dREMLfit)* head-motion parameters, movement spike regressors (frame wise displacement above 0.3) and the mean signal of cerebro-spinal fluid (CSF), as well as accounting for the temporal autoregression (*ARMA-1*), which typically reflects artifacts of physiological origin (Bright et al., 2017). The pre-processed data were then non-linearly transformed *(3dNwarpApply)* into the Montreal Neurological Institute (MNI152) coordinate system and resampled to a 2 mm iso-voxel resolution. Finally, to reduce computational effort in the subsequent steps, a gray matter spatial mask was also applied (p > 0.10 in the tissue probability map of the ICBM 2009c atlas; Fonov et al., 2009).

Brain regions associated with the occurrence of sleep slow waves were identified through a voxel-wise regression of BOLD time-series. Specifically, for each subject, an EEG-based regressor was built, in which each slow wave was modelled as a square wave with onset-time corresponding to the timing of the first zero-crossing of the negative half-wave, height equal to the absolute value of the maximum negative peak (*A*), and duration corresponding to the length of the descending phase of the wave (from first zero-crossing to negative peak; *d*_1_). The obtained regressor was then convoluted with a standard gamma hemodynamic response function (p = 8.6, q= 0.547) and down-sampled to the BOLD time-series sampling rate (0.5 Hz, TR = 2 s). The regressor and each BOLD time-series were forced respectively to zero and the baseline value in correspondence of all artifactual and wakefulness epochs (censored intervals). At single-subject level, beta-values of all cortical voxels were converted into z-scores calculated with respect to a null distribution obtained by re-computing the regression analysis on the same BOLD time-series after shuffling the timing of individual slow waves (intra-voxel and - subject permutations; nPerm = 1000). Importantly, the number and amplitude of slow waves was kept constant across original and shuffled regressors (i.e., relocation of slow waves within censored intervals was prevented). A group-level one-sample t-test was then performed at each voxel to assess statistical significance *(3dttest)*. The significance threshold was set to q < 0.01 after FDR correction for multiple comparisons (False Discovery Rate; Benjamini and Hochberg, 1995). A minimum, arbitrary cluster-size threshold corresponding to 50 voxels was also applied.

### Thalamic and cerebellar involvement in sleep slow waves

Specific analyses were performed to characterize the anatomical and functional nature of thalamic and cerebellar portions recruited during the occurrence of human sleep slow waves. First, anatomical atlases were used to determine the relative distribution (percentage) of activated voxels with respect to thalamic (probabilistic atlas based on diffusion-weighted imaging; Najdenovska et al., 2018) and cerebellar (SUIT atlas; Diedrichsen et al., 2011, 2009) subdivisions. Then, thalamic and cerebellar connectivity maps were generated and used to determine the preferential functional connectivity of activated voxels with respect to the seven canonical cortical networks defined on the basis of cortical intrinsic functional connectivity (Yeo et al., 2011): visual, somatomotor, dorsal attention, ventral attention, limbic, fronto-parietal and default mode. This analysis was performed using rs-fMRI data collected during wakefulness in an independent sample of 28 subjects (see Participants section; T2*-weighted gradient-echo echoplanar sequences with 35 axial contiguous slices, TR = 2000 ms; TE = 35 ms; FA = 80°; voxel size: 3×3×4 mm; 230 volumes per run). In particular, for each subject, we first computed the average BOLD-signal within 200 cortical ROIs, as defined in Schaefer et al. (2018). Then, the resting state signal from the 200 ROIs was further averaged according to the seven cortical networks, separately for each brain hemisphere (thus leading to a total of 14 large ROIs). For what concerns the thalamus, we then evaluated the partial correlation between the thalamic signal of each voxel and each homolateral cortical functional network while removing signal variance from all the left-out functional networks (Hwang et al, 2017). Regarding the cerebellum, we applied a similar procedure with two important differences. First, we measured the partial correlation between each cerebellar voxel and the contralateral cortical functional networks. Second, as described in previous work (Buckner et al., 2011), the mean signal of neocortical voxels located in the proximity of the cerebellum (up to 6 mm) was additionally included as variable of no interest in the partial correlation procedure. This procedure ensured that the putative somatomotor region of the cerebellar anterior lobe remained relatively unaffected by the possible ‘leakage’ of BOLD-signal from the spatially close visual cortex. The described partial correlation procedure led to the generation of a connectivity matrix in each subject. A group-level matrix was obtained by computing the median across individual matrices. Finally, a winner-takes-all approach was applied to assign the local ‘preferential connectivity’ of each voxel (Hwang et al., 2017). This procedure allowed to generate functional connectivity maps of the thalamus and the cerebellum similar to those reported in Hwang et al. (2017) and Buckner et al. (2011), respectively. Then, for both the thalamus and the cerebellum, we computed the proportion of voxels connected with each of the seven networks that resulted included in the significantly activated regional clusters (see Figures 4 and 5).

### Cortical involvement in sleep slow waves

At cortical level, the sleep slow waves are known to propagate through white matter anatomical pathways between brain areas (Avvenuti et al., 2020; Massimini et al., 2004; Murphy et al., 2009). Thus, here additional analyses were performed to determine whether the specific changes in hemodynamic activity observed in association with sleep slow waves remain confined within specific cortical sites or may propagate with some delay to other cortical areas following the spreading of electrophysiological slow waves. To this aim, we used a crosscorrelation procedure to compute the similarity and relative lag of activity in different brain areas and a *‘seed’* region chosen as a reference template. Specifically, the left somatomotor cortex, as defined based on the regression analysis, was here selected as *seed* region (though similar results were obtained using different *seed* areas, such as the right somatomotor cortex or the right visual cortex; *data not shown).* Then, the *seed* time-series was compared with the average BOLD-signals computed for each of the 200 ROIs of the Schaefer atlas (Schaefer et al., 2018). In particular, BOLD time-series in a timewindow ranging from −4 s to +16 s with respect to slow-wave onset were first averaged across voxels belonging to each ROI, then across slow-waves and finally across subjects. The time-window of interest was selected in order to mainly include the negative portion of the hemodynamic response (Figure 6). Such a choice was made for two reasons. First, the negative deflection is larger with respect to the slower positive deflection observed before slow-wave onset. Second, the negative deflection may be expected to more likely reflect a consequence of the neuronal silence that is associated with the *down-state* of sleep slow waves (Fultz et al., 2019). Of note, time-series included in this analysis were up-sampled to 256 Hz (linear interpolation), thus matching the EEG sampling rate, in order to improve the alignment with the timing of slow-wave onset. The cross-correlation function between the signal profile of the template and the mean time-series of each ROI was computed to estimate the relative time-lag that allowed to maximize the similarity between each pair of time-series. In addition, the magnitude of the corresponding peak in the cross-correlation function was used to evaluate the statistical significance of the similarity between examined signals. To this aim, the above-mentioned procedure was repeated following a random shuffling of the onset-timing of individual slow waves in each subject (nPerm = 1000), thus eventually obtaining a null-distribution of similarity values. ROIs for which the maximum similarity values were significantly greater than those in the null-distribution (p < 0.05) were used to build a propagation delay-map in which all regional lag values were re-scaled to the minimum delay-value in the map (t = 0 s; Figure 7).

In order to determine whether cortical delays in hemodynamic changes may reflect the actual propagation of electrophysiological slow waves we directly compared the fMRI-based delay-map with a delay-map of EEG slow-wave propagation in source space. This map was obtained from the analysis of NREM-sleep hd-EEG recordings (256 electrodes, EGI-Philips; 500Hz sampling frequency) obtained in a distinct set of 12 healthy adult volunteers (see Participants section). For each participant, the first 30 minutes of N2/N3 sleep were extracted and slow waves were automatically detected using the same algorithm described above. Then, the EEG signal was band-pass filtered between 0.5 and 4.0 Hz, and 800-ms-long data segments centered on the slow wave negative peak were extracted and source-modeled using the GeoSource 3.0 software (EGI-Philips), as described in previous work (Bernardi et al., 2019a, 2019b). In brief, a four-shell head model based on the MNI atlas and subject-specific co-registered sets of electrode positions (obtained using the Geodesic Photogrammetry System, EGI-Philips) were used to construct the forward model. The inverse matrix was computed using the standardized low-resolution brain electromagnetic tomography constraint (sLORETA; Tikhonov regularization = 10^-2^). The source space was restricted to 2447 dipoles distributed over 7 mm^3^ cortical voxels. For each slow wave, the propagation pattern was determined using an approach similar to the one described by Murphy and colleagues (Murphy et al., 2009). Specifically, for each slow wave, we defined a time-window of 100 ms centered on the timing of the maximum negative peak of the slow wave detected at scalp level. Next, for each voxel, we computed the timing of any local maxima that occurred during the time window of interest. For each voxel, after discarding secondary peaks (defined as peaks with magnitude lower that the 75% of the maximum peak across voxels), we selected the maxima that occurred most closely to the reference peak. The relative timing of these local maxima were used to create a preliminary propagation delay-map. A spatio-temporal clusterization procedure was applied with the aim of excluding potential propagation gaps (Avvenuti et al., 2020): local peaks of two spatial neighbor voxels had to be separated by less than 10 ms in order to be considered as part of the same propagation cluster. The propagation cluster including the largest signal peak was then identified and used to extract the final delay-map. The minimum observed delay, corresponding to the slow wave origin, was set to zero. Delay-maps obtained for each wave and subject were averaged in order to obtain a group-level delay-map. In order to allow a comparison with the BOLD delay-map, the EEG-based delay-map was resampled to a 2 mm iso-voxel resolution, matching the spatial resolution of the fMRI dataset, and mean delay values were computed for each of the 200 ROIs of the Schaefer atlas (Schaefer et al., 2018). The Pearson’s correlation coefficient was eventually used to quantify the similarity between fMRI and EEG delay-maps.

### Relationship between slow-wave amplitude and BOLD-signal variations

Finally, we investigated the possible relationship between the amplitude of EEG slow waves and corresponding variations in hemodynamic responses within cortical and subcortical structures. To this aim, we first performed a time-wise correlation analysis between slow wave amplitude and point-by-point (up-sampled) BOLD-signal changes for each of the significant clusters obtained from the regression analysis. In addition, a trend analysis was performed in the same areas to determine whether a relationship exists between slow wave amplitude and the delay of the regional BOLD response. Specifically, the delay of each BOLD profile was defined as the timing (relative to slow-wave onset) of the maximum positive or negative peak in subcortical and cortical regions, respectively. For this analysis, EEG slow waves were divided in three percentile classes (*A_1_*: 0-33; *A_2_*: 33-66; *A_3_*: 66-100) based on their negative amplitude, and the mean delay was computed for each class. The potential trend of the mean delay as a function of EEG amplitude was then quantified with a regression analysis. Significance of the results for both the correlation and the trend analyses was tested using a within-subject permutation-based approach (nPerm = 10000), in which the same procedures were applied after randomly shuffling the correspondence between EEG amplitudes and fMRI-derived values. Obtained p-values were separately evaluated for the positive and negative tails of the distribution, combined across subjects using the Fisher’s method (Brown, 1975) and FDR adjusted to correct for multiple comparisons.

### Data availability

All relevant cortical and subcortical maps, including main results and parcellations, will be made freely available in a public data repository upon publication.

## Acknowledgments

This work was supported by intramural funds from the IMT School for Advanced Studies Lucca (to G.B., M.B., G.H., A.L.), the Swiss National Science Foundation (Ambizione Grant PZ00P3_173955 to F.S.), the Divesa Foundation Switzerland (to F.S.), the Pierre-Mercier Foundation for Science (to F.S.), the Bourse Pro-Femme of the University of Lausanne (to F.S.), the Foundation for the University of Lausanne (to F.S and G.B.), an International Brain Research Organization Pan-European Regional Committee short-term postdoctoral fellowship (to G.B.), and a grant “Dipartimenti di Eccellenza 2018-2022”, MIUR-Italy, to the Department of Biomedical, Metabolic and Neural Sciences (S.M. and F.B.).

## Author Contributions

*Conceptualization:* G.B., F.S., G.H., F.B., E.R.; *Data acquisition:* M.B., G.H., A.L., V.F., S.M., D.B., F.B., G.B.; *Data analysis:* M.B., G.H., A.L., A.F., G.B.; *Visualization:* M.B., G.H, G.B.*; Results interpretation:* G.B., M.B., F.S.; *Wrote the manuscript:* G.B., M.B., G.H.; *Revised and edited the manuscript*: All authors.

## References

Allen PJ, Josephs O, Turner R. 2000. A method for removing imaging artifact from continuous EEG recorded during functional MRI. Neuroimage 12:230–239. doi:10.1006/nimg.2000.0599

Amzica F, Steriade M. 1995. Short-and long-range neuronal synchronization of the slow (<1 Hz) cortical oscillation. J Neurophysiol 73:20–38. doi:10.1152/jn.1995.73.1.20

Andrillon T, Kouider S. 2020. The vigilant sleeper: neural mechanisms of sensory (de)coupling during sleep. Curr Opin Physiol 15:47–59. doi:10.1016/j.cophys.2019.12.002

Avvenuti G, Handjaras G, Betta M, Cataldi J, Imperatori LS, Lattanzi S, Riedner BA, Pietrini P, Ricciardi E, Tononi G, Siclari F, Polonara G, Fabri M, Silvestrini M, Bellesi M, Bernardi G. 2020. Integrity of corpus callosum is essential for the cross-hemispheric propagation of sleep slow waves: a high-density EEG study in split-brain patients. J Neurosci In Press.

Benjamini Y, Hochberg Y. 1995. Controlling the false discovery rate: a practical and powerful approach to multiple testing. J R Stat Soc Ser B 289–300.

Bernardi G, Betta M, Cataldi J, Leo A, Haba-Rubio J, Heinzer R, Cirelli C, Tononi G, Pietrini P, Ricciardi E, Siclari F. 2019a. Visual imagery and visual perception induce similar changes in occipital slow waves of sleep. J Neurophysiol 121:2140–2152. doi:10.1152/jn.00085.2019

Bernardi G, Betta M, Ricciardi E, Pietrini P, Tononi G, Siclari F. 2019b. Regional delta waves in human rapid eye movement sleep. J Neurosci 39:2686–2697. doi:10.1523/JNEUROSCI.2298-18.2019

Bernardi G, Siclari F, Handjaras G, Riedner BA, Tononi G. 2018. Local and Widespread Slow Waves in Stable NREM Sleep: Evidence for Distinct Regulation Mechanisms. Front Hum Neurosci 12:1–13. doi:10.3389/fnhum.2018.00248

Bright MG, Tench CR, Murphy K. 2017. Potential pitfalls when denoising resting state fMRI data using nuisance regression. Neuroimage 154:159–168. doi:10.1016/j.neuroimage.2016.12.027

Brown MB. 1975. A Method for Combining Non-Independent, One-Sided Tests of Significance. Biometrics 31:987. doi:10.2307/2529826

Brown RE, Basheer R, McKenna JT, Strecker RE, McCarley RW. 2012. Control of Sleep and Wakefulness. Physiol Rev 92:1087–1187. doi:10.1152/physrev.00032.2011

Buchmann A, Kurth S, Ringli M, Geiger A, Jenni OG, Huber R. 2011. Anatomical markers of sleep slow wave activity derived from structural magnetic resonance images. J Sleep Res 20:506–513. doi:10.1111/j.1365-2869.2011.00916.x

Buckner RL, Krienen FM, Castellanos A, Diaz JC, Thomas Yeo BT. 2011. The organization of the human cerebellum estimated by intrinsic functional connectivity. J Neurophysiol 106:2322–2345. doi:10.1152/jn.00339.2011

Canto CB, Onuki Y, Bruinsma B, van der Werf YD, De Zeeuw CI. 2017. The Sleeping Cerebellum. Trends Neurosci 40:309–323. doi:10.1016/j.tins.2017.03.001

Colavito V, Tesoriero C, Wirtu AT, Grassi-Zucconi G, Bentivoglio M. 2015. Limbic thalamus and state-dependent behavior: The paraventricular nucleus of the thalamic midline as a node in circadian timing and sleep/wake-regulatory networks. Neurosci Biobehav Rev 54:3–17. doi:10.1016/j.neubiorev.2014.11.021

Contreras D, Steriade M. 1995. Cellular basis of EEG slow rhythms: A study of dynamic corticothalamic relationships. J Neurosci 15:604–622. doi:10.1523/jneurosci.15-01-00604.1995

Cox R, Van Driel J, De Boer M, Talamini LM. 2014. Slow oscillations during sleep coordinate interregional communication in cortical networks. J Neurosci 34:16890–16901. doi:10.1523/JNEUROSCI.1953-14.2014

Cox RW. 2012. AFNI: What a long strange trip it’s been. Neuroimage 62:743–747. doi:10.1016/j.neuroimage.2011.08.056

Cox RW. 1996. AFNI: software for analysis and visualization of functional magnetic resonance neuroimages. Comput Biomed Res 29:162–73. doi:10.1006/cbmr.1996.0014

Crunelli V, Hughes SW. 2010. The slow (1 Hz) rhythm of non-REM sleep: A dialogue between three cardinal oscillators. Nat Neurosci 13:9–17. doi:10.1038/nn.2445

Crunelli V, Larincz ML, Connelly WM, David F, Hughes SW, Lambert RC, Leresche N, Errington AC. 2018. Dual function of thalamic low-vigilance state oscillations: Rhythm-regulation and plasticity. Nat Rev Neurosci 19:107–118. doi:10.1038/nrn.2017.151

Dang-Vu TT, Schabus M, Desseilles M, Albouy G, Boly M, Darsaud A, Gais S, Rauchs G, Sterpenich V, Vandewalle G, Carrier J, Moonen G, Balteau E, Degueldre C, Luxen A, Phillips C, Maquet P. 2008. Spontaneous neural activity during human slow wave sleep. Proc Natl Acad Sci USA 105:15160–5. doi:10.1073/pnas.0801819105

David F, Schmiedt JT, Taylor HL, Orban G, Di Giovanni G, Uebele VN, Renger JJ, Lambert RC, Leresche N, Crunelli V. 2013. Essential thalamic contribution to slow waves of natural sleep. J Neurosci 33:19599–19610. doi:10.1523/JNEUROSCI.3169-13.2013

De Gennaro L, Gorgoni M, Reda F, Lauri G, Truglia I, Cordone S, Scarpelli S, Mangiaruga A, D’Atri A, Lacidogna G, Ferrara M, Marra C, Rossini PM. 2017. The Fall of Sleep K-Complex in Alzheimer Disease. Sci Rep 7. doi:10.1038/srep39688

Delorme A, Makeig S. 2004. EEGLAB: an open source toolbox for analysis of single-trial EEG dynamics including independent component analysis. J Neurosci Methods 134:9–21.

Diedrichsen J, Balsters JH, Flavell J, Cussans E, Ramnani N. 2009. A probabilistic MR atlas of the human cerebellum. Neuroimage 46:39–46. doi:10.1016/j.neuroimage.2009.01.045

Diedrichsen J, Maderwald S, Küper M, Thürling M, Rabe K, Gizewski ER, Ladd ME, Timmann D. 2011. Imaging the deep cerebellar nuclei: A probabilistic atlas and normalization procedure. Neuroimage 54:1786–1794. doi:10.1016/j.neuroimage.2010.10.035

Diekelmann S, Born J. 2010. The memory function of sleep. Nat Rev Neurosci 11:114–126.

Eschenko O, Magri C, Panzeri S, Sara SJ. 2012. Noradrenergic neurons of the locus coeruleus are phase locked to cortical up-down states during sleep. Cereb Cortex 22:426–435. doi:10.1093/cercor/bhr121

Fonov V, Evans A, McKinstry R, Almli C, Collins D. 2009. Unbiased nonlinear average age-appropriate brain templates from birth to adulthood. Neuroimage 47:S102. doi:10.1016/s1053-8119(09)70884-5

Fultz NE, Bonmassar G, Setsompop K, Stickgold RA, Rosen BR, Polimeni JR, Lewis LD. 2019. Coupled electrophysiological, hemodynamic, and cerebrospinal fluid oscillations in human sleep. Science (80-) 366:628–631.

Funk CM, Honjoh S, Rodriguez A V, Cirelli C, Tononi G. 2016. Local Slow Waves in Superficial Layers of Primary Cortical Areas during REM Sleep. Curr Biol 26:396–403.

Gaspar P, Berger B, Febvret A, Vigny A, Henry JP. 1989. Catecholamine innervation of the human cerebral cortex as revealed by comparative immunohistochemistry of tyrosine hydroxylase and dopamine-beta-hydroxylase. J Comp Neurol 279:249–271. doi:10.1002/cne.902790208

Gent TC, Bandarabadi M, Herrera CG, Adamantidis AR. 2018a. Thalamic dual control of sleep and wakefulness. Nat Neurosci 21:974–984. doi:10.1038/s41593-018-0164-7

Gent TC, Bassetti C LA, Adamantidis AR. 2018b. Sleep-wake control and the thalamus. Curr Opin Neurobiol 52:188–197. doi:10.1016/j.conb.2018.08.002

Hablitz LM, Vinitsky HS, Sun Q, Stæger FF, Sigurdsson B, Mortensen KN, Lilius TO, Nedergaard M. 2019. Increased glymphatic influx is correlated with high EEG delta power and low heart rate in mice under anesthesia. Sci Adv 5:eaav5447. doi:10.1126/sciadv.aav5447

Honjoh S, Sasai S, Schiereck SS, Nagai H, Tononi G, Cirelli C. 2018. Regulation of cortical activity and arousal by the matrix cells of the ventromedial thalamic nucleus. Nat Commun 9. doi:10.1038/s41467-018-04497-x

Hwang K, Bertolero MA, Liu WB, D’Esposito M. 2017. The human thalamus is an integrative hub for functional brain networks. J Neurosci 37:5594–5607. doi:10.1523/JNEUROSCI.0067-17.2017

Iannetti GD, Niazy RK, Wise RG, Jezzard P, Brooks JCW, Zambreanu L, Vennart W, Matthews PM, Tracey I. 2005. Simultaneous recording of laser-evoked brain potentials and continuous, high-field functional magnetic resonance imaging in humans. Neuroimage 28:708–719. doi:10.1016/j.neuroimage.2005.06.060

Iber C, Ancoli-Israel S A C. 2007. The AASM manural for the scoring of sleep and associated events: Rules, terminology and technical specifications, American Academy of Sleep Medicine.

Jahnke K, von Wegner F, Morzelewski A, Borisov S, Maischein M, Steinmetz H, Laufs H. 2012. To wake or not to wake? The two-sided nature of the human K-complex. Neuroimage 59:1631–1638. doi:10.1016/J.NEUROIMAGE.2011.09.013

Javoy-Agid F, Scatton B, Ruberg M, L’heureux R, Cervera P, Raisman R, Maloteaux J-M, Beck H, Agid Y. 1989. Distribution of monoaminergic, cholinergic, and GABAergic markers in the human cerebral cortex. Neuroscience 29:251–259. doi:10.1016/0306-4522(89)90055-9

Laurino M, Menicucci D, Piarulli A, Mastorci F, Bedini R, Allegrini P, Gemignani A. 2014. Disentangling different functional roles of evoked K-complex components: mapping the sleeping brain while quenching sensory processing. Neuroimage 86:433–445.

Laurino M, Piarulli A, Menicucci D, Gemignani A. 2019. Local Gamma Activity During Non-REM Sleep in the Context of Sensory Evoked K-Complexes. Front Neurosci 13. doi:10.3389/fnins.2019.01094

Lemieux M, Chen JY, Lonjers P, Bazhenov M, Timofeev I. 2014. The impact of cortical deafferentation on the neocortical slow oscillation. J Neurosci 34:5689–5703. doi:10.1523/JNEUROSCI.1156-13.2014

Lewis DA, Morrison JH. 1989. Noradrenergic innervation of monkey prefrontal cortex: A dopamine-β-hydroxylase immunohistochemical study. J Comp Neurol 282:317–330. doi:10.1002/cne.902820302

Lőrincz ML, Gunner D, Bao Y, Connelly WM, Isaac JTR, Hughes SW, Crunelli V. 2015. A distinct class of slow (~0.2-2 Hz) intrinsically bursting layer 5 pyramidal neurons determines UP/DOWN state dynamics in the neocortex. J Neurosci 35:5442–5458. doi:10.1523/JNEUROSCI.3603-14.2015

Mander BA, Marks SM, Vogel JW, Rao V, Lu B, Saletin JM, Ancoli-Israel S, Jagust WJ, Walker MP. 2015. β-amyloid disrupts human NREM slow waves and related hippocampus-dependent memory consolidation. Nat Neurosci 18:1051–1057. doi:10.1038/nn.4035

Massimini M, Huber R, Ferrarelli F, Hill S, Tononi G. 2004. The Sleep Slow Oscillation as a Traveling Wave. J Neurosci 24:6862–6870. doi:10.1523/JNEUROSCI.1318-04.2004

Mena-segovia J, Sims HM, Magill PJ, Bolam JP. 2008. Cholinergic brainstem neurons modulate cortical gamma activity during slow oscillations. J Physiol 586:2947–2960. doi:10.1113/jphysiol.2008.153874

Menicucci D, Piarulli A, Debarnot U, d’Ascanio P, Landi A, Gemignani A. 2009. Functional structure of spontaneous sleep slow oscillation activity in humans. PLoS One 4:e7601.

Mensen A, Riedner B, Tononi G. 2016. Optimizing detection and analysis of slow waves in sleep EEG. J Neurosci Methods 274:1–12.

Mitra A, Snyder AZ, Tagliazucchi E, Laufs H, Raichle ME. 2015. Propagated infra-slow intrinsic brain activity reorganizes across wake and slow wave sleep. Elife 4. doi:10.7554/elife.10781

Miyamoto D, Hirai D, Murayama M. 2017. The roles of cortical slow waves in synaptic plasticity and memory consolidation. Front Neural Circuits 11. doi:10.3389/fncir.2017.00092

Moruzzi G, Magoun HWW. 1949. Brain stem reticular formation and activation of the EEG. Electroencephalogr Clin Neurophysiol 1:455–473. doi:10.1016/0013-4694(49)90219-9

Mountcastle VB, Poggio GF. 1974. Structural organization and general physiology of thalamotelencephalic systemsMedical Physiology. CV Mosby Co St. Louis. pp. 227–253.

Mullinger K, Bowtell R. 2010. Combining EEG and fMRI. Methods Mol Biol 711:303–326. doi:10.1007/978-1-61737-992-5_15

Murphy M, Riedner BA, Huber R, Massimini M, Ferrarelli F, Tononi G. 2009. Source modeling sleep slow waves. Proc Natl Acad Sci 106:1608–1613.

Najdenovska E, Alemán-Gómez Y, Battistella G, Descoteaux M, Hagmann P, Jacquemont S, Maeder P, Thiran IP, Fornari E, Cuadra MB. 2018. In-vivo probabilistic atlas of human thalamic nuclei based on diffusion-weighted magnetic resonance imaging. Sci Data 5. doi:10.1038/sdata.2018.270

Neske GT. 2016. The Slow Oscillation in Cortical and Thalamic Networks: Mechanisms and Functions. Front Neural Circuits 9. doi:10.3389/fncir.2015.00088

Niazy RK, Beckmann CF, Iannetti GD, Brady JM, Smith SM. 2005. Removal of FMRI environment artifacts from EEG data using optimal basis sets. Neuroimage 28:720–737. doi:10.1016/j.neuroimage.2005.06.067

Piayantoni G, Poil S-S, Linkenkaer-Hansen K, Verweij IM, Ramautar JR, Van Someren EJW, Van Der Werf YD. 2013. Individual Differences in White Matter Diffusion Affect Sleep Oscillations. J Neurosci 33:227–233. doi:10.1523/jneurosci.2030-12.2013

Poulet JFA, Fernandez LMJ, Crochet S, Petersen CCH. 2012. Thalamic control of cortical states. Nat Neurosci 15:370–372. doi:10.1038/nn.3035

Riedner BA, Hulse BK, Murphy MJ, Ferrarelli F, Tononi G. 2011. Temporal dynamics of cortical sources underlying spontaneous and peripherally evoked slow waves. Prog Brain Res 193:201.

Riedner BA, Vyazovskiy V V, Huber R, Massimini M, Esser S, Murphy M, Tononi G. 2007. Sleep homeostasis and cortical synchronization: III. A high-density EEG study of sleep slow waves in humans. Sleep 30:1643.

Roš H, Sachdev RNS, Yu Y, Šestan N, McCormick DA. 2009. Neocortical networks entrain neuronal circuits in cerebellar cortex. J Neurosci 29:10309–10320. doi:10.1523/JNEUROSCI.2327-09.2009

Rowland NC, Goldberg JA, Jaeger D. 2010. Cortico-cerebellar coherence and causal connectivity during slow-wave activity. Neuroscience 166:698–711. doi:10.1016/j.neuroscience.2009.12.048

Sanchez-Vives M V., McCormick DA. 2000. Cellular and network mechanisms of rhytmic recurrent activity in neocortex. Nat Neurosci 3:1027–1034. doi:10.1038/79848

Schaefer A, Kong R, Gordon EM, Laumann TO, Zuo X-N, Holmes AJ, Eickhoff SB, Yeo BTT. 2018. Local-Global Parcellation of the Human Cerebral Cortex from Intrinsic Functional Connectivity MRI. Cereb Cortex 28:3095–3114. doi:10.1093/cercor/bhx179

Schweimer J V., Mallet N, Sharp T, Ungless MA. 2011. Spike-timing relationship of neurochemically-identified dorsal raphe neurons during cortical slow oscillations. Neuroscience 196:115–123. doi:10.1016/j.neuroscience.2011.08.072

Sheroziya M, Timofeev I. 2014. Global intracellular slow-wave dynamics of the thalamocortical system. J Neurosci 34:8875–8893. doi:10.1523/JNEUROSCI.4460-13.2014

Siclari F, Bernardi G, Riedner BA, LaRocque JJ, Benca RM, Tononi G. 2014. Two Distinct Synchronization Processes in the Transition to Sleep: A High-Density Electroencephalographic Study. Sleep 37:1621–37. doi:10.5665/sleep.4070

Slézia A, Hangya B, Ulbert I, Acsády L. 2011. Phase advancement and nucleus-specific timing of thalamocortical activity during slow cortical oscillation. J Neurosci 31:607–617. doi:10.1523/JNEUROSCI.3375-10.2011

Steriade M, Aposto V, Oakson G. 1971a. Clustered firing in the cerebello-thalamic pathway during synchronized sleep. Brain Res 26:425–432. doi:10.1016/S0006-8993(71)80020-3

Steriade M, Apostol V, Oakson G. 1971b. Control of unitary activities in cerebellothalamic pathway during wakefulness and synchronized sleep. J Neurophysiol 34:389–413. doi:10.1152/jn.1971.34.3.389

Steriade M, Glenn LL. 1982. Neocortical and caudate projections of intralaminar thalamic neurons and their synaptic excitation from midbrain reticular core. J Neurophysiol 48:352–371. doi:10.1152/jn.1982.48.2.352

Steriade M, Nunez A, Amzica F. 1993a. A novel slow (< 1 Hz) oscillation of neocortical neurons in vivo: Depolarizing and hyperpolarizing components. J Neurosci 13:3252–3265. doi:10.1523/jneurosci.13-08-03252.1993

Steriade M, Nunez A, Amzica F. 1993b. Intracellular analysis of relations between the slow (<1 Hz) neocortical oscillation and other sleep rhythms of the electroencephalogram. J Neurosci 13:3266–3283. doi:10.1523/jneurosci.13-08-03266.1993

Thomas Yeo BT, Krienen FM, Sepulcre J, Sabuncu MR, Lashkari D, Hollinshead M, Roffman JL, Smoller JW, Zöllei L, Polimeni JR, Fisch B, Liu H, Buckner RL. 2011. The organization of the human cerebral cortex estimated by intrinsic functional connectivity. J Neurophysiol 106:1125–1165. doi:10.1152/jn.00338.2011

Timofeev I, Chauvette S. 2017. Sleep slow oscillation and plasticity. Curr Opin Neurobiol 44:116–126. doi:10.1016/j.conb.2017.03.019

Timofeev I, Grenier F, Bazhenov M, Sejnowski TJ, Steriade M, Timofeev, I; Grenier, F; Bazhenov, M; Sejnowski, TJ, Steriade M. 2000. Origin of slow cortical oscillations in deafferented cortical slabs. Cereb Cortex 10:1185–1199.

Timofeev I, Steriade M. 1996. Low-frequency rhythms in the thalamus of intact-cortex and decorticated cats. J Neurophysiol 76:4152–4168. doi:10.1152/jn.1996.76.6.4152

Tononi G, Cirelli C. 2014. Sleep and the price of plasticity: from synaptic and cellular homeostasis to memory consolidation and integration. Neuron 81:12–34. doi:10.1016/j.neuron.2013.12.025

Ushimaru M, Kawaguchi Y. 2015. Temporal structure of neuronal activity among cortical neuron subtypes during slow oscillations in anesthetized rats. J Neurosci 35:11988–12001. doi:10.1523/JNEUROSCI.5074-14.2015

Vertes RP, Linley SB, Hoover WB. 2015. Limbic circuitry of the midline thalamus. Neurosci Biobehav Rev 54:89–107. doi:10.1016/j.neubiorev.2015.01.014

Xie L, Kang H, Xu Q, Chen MJ, Liao Y, Thiyagarajan M, O’Donnell J, Christensen DJ, Nicholson C, Iliff JJ. 2013. Sleep drives metabolite clearance from the adult brain. Science (80-) 342:373–377.

