## Supplementary for "Cortical and subcortical hemodynamic changes during human sleep slow waves"


**Figure S1**

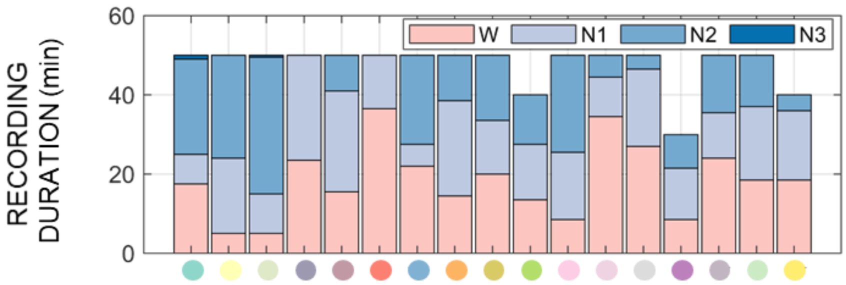


| STG | TOT |
|---|---|
| W | 312.5 |
| N1 | 266.0 |
| N2 | 230.0 |
| N3 | 1.5 |

*Figure S1. Distribution of sleep stages in each participant. Each column represents a different subject. Three participants interrupted the scan session before completion of the five scheduled EEG-fMRI runs because they felt unable to fall asleep again. The table on the right shows the total amount of each sleep stage (stg) in minutes.*

**Figure S2**

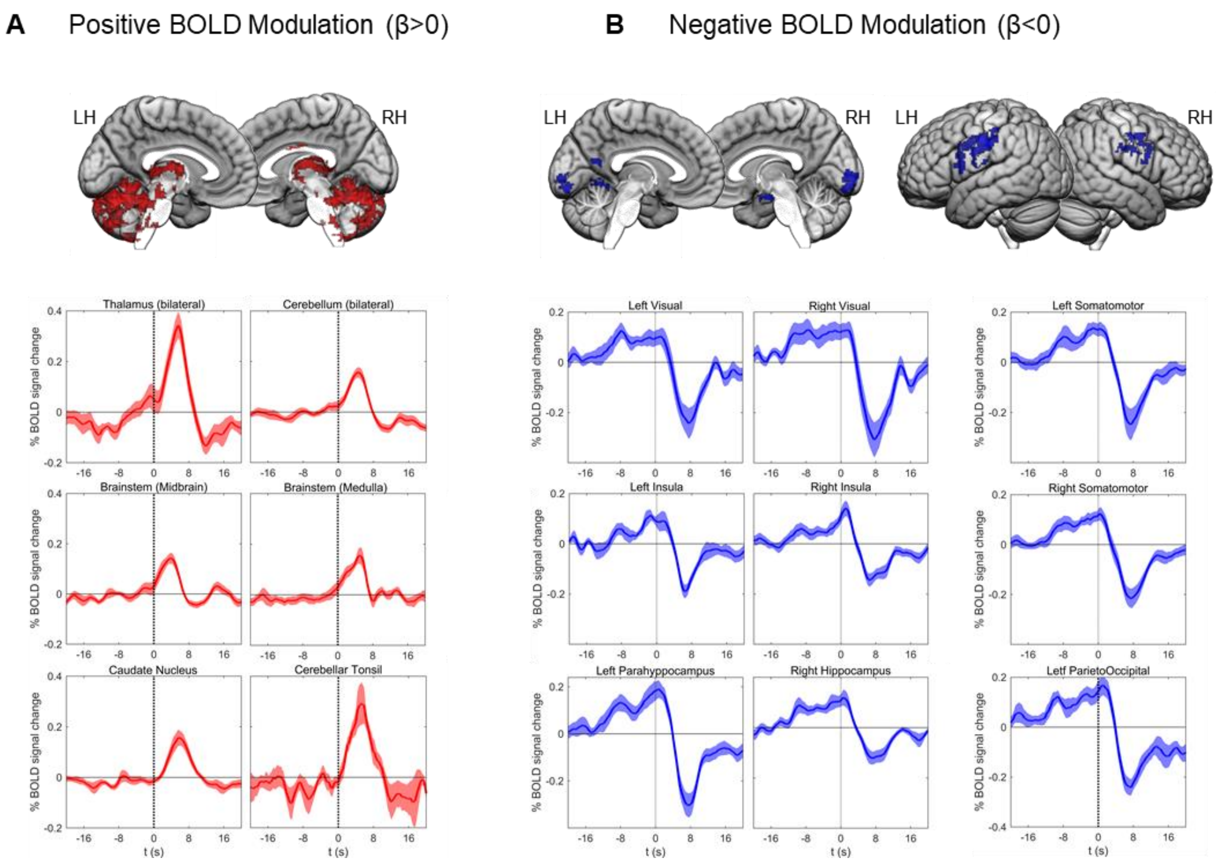


*Figure S2. Hemodynamic changes associated with sleep slow waves. Mean BOLD-signals (up-sampled to the EEG sampling rate) and relative standard errors for all identified significant clusters (top row). Time t = 0 s corresponds to slow-wave onset. Plots on the left (A; in red) show areas associated with a predominant positive BOLD-signal change, while plots on the right (B; in blue) show areas associated with a predominant negative BOLD-signal change.*

**Figure S3**

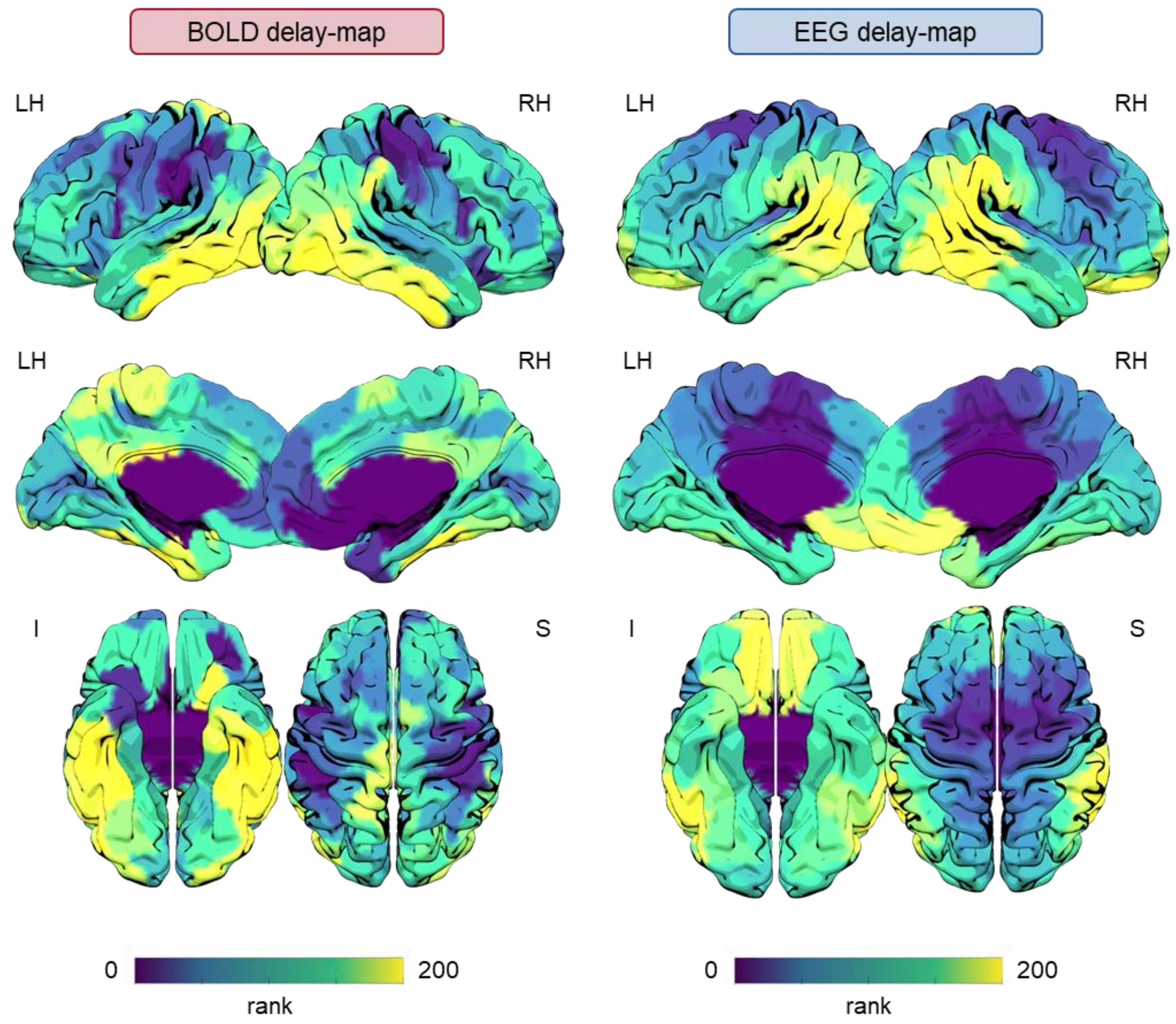


*Figure S3. Hemodynamic and electrophysiological signal propagation during sleep slow waves. Rank-based cortical delay-maps. In order to create these images, the 200 ROIs of the Schaefer atlas were ranked based on their relative mean delay (see main text for details regarding how delays were computed for BOLD- and EEG-signals). Brain surface plots were generated using the Surf Ice software (https://www.nitrc.org/projects/surfice/). Here the yellow color indicates a high rank (high delay), while the dark-blue color indicates a low rank (low delay).*

**Figure S4**

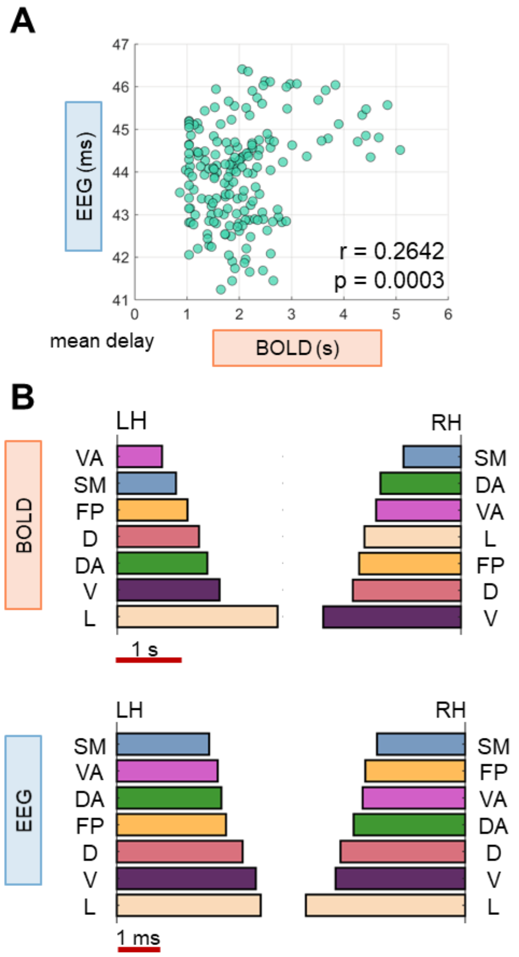


*Figure S4. Coupled hemodynamic and electrophysiological signal propagation during sleep slow waves. (A) Correlation between EEG- and BOLD-signal delays for 182 ROIs of the Schaefer atlas that showed a significant cross-correlation with the seed region in the computation of the BOLD delay-map. (B) Mean propagation delays computed for the seven canonical networks, separately for the left and the right hemisphere. SM: somatomotor, VA: ventral attention, FP: fronto-parietal, DA: dorsal attention, D: default mode, V: visual, L: limbic.*

**Figure S5**

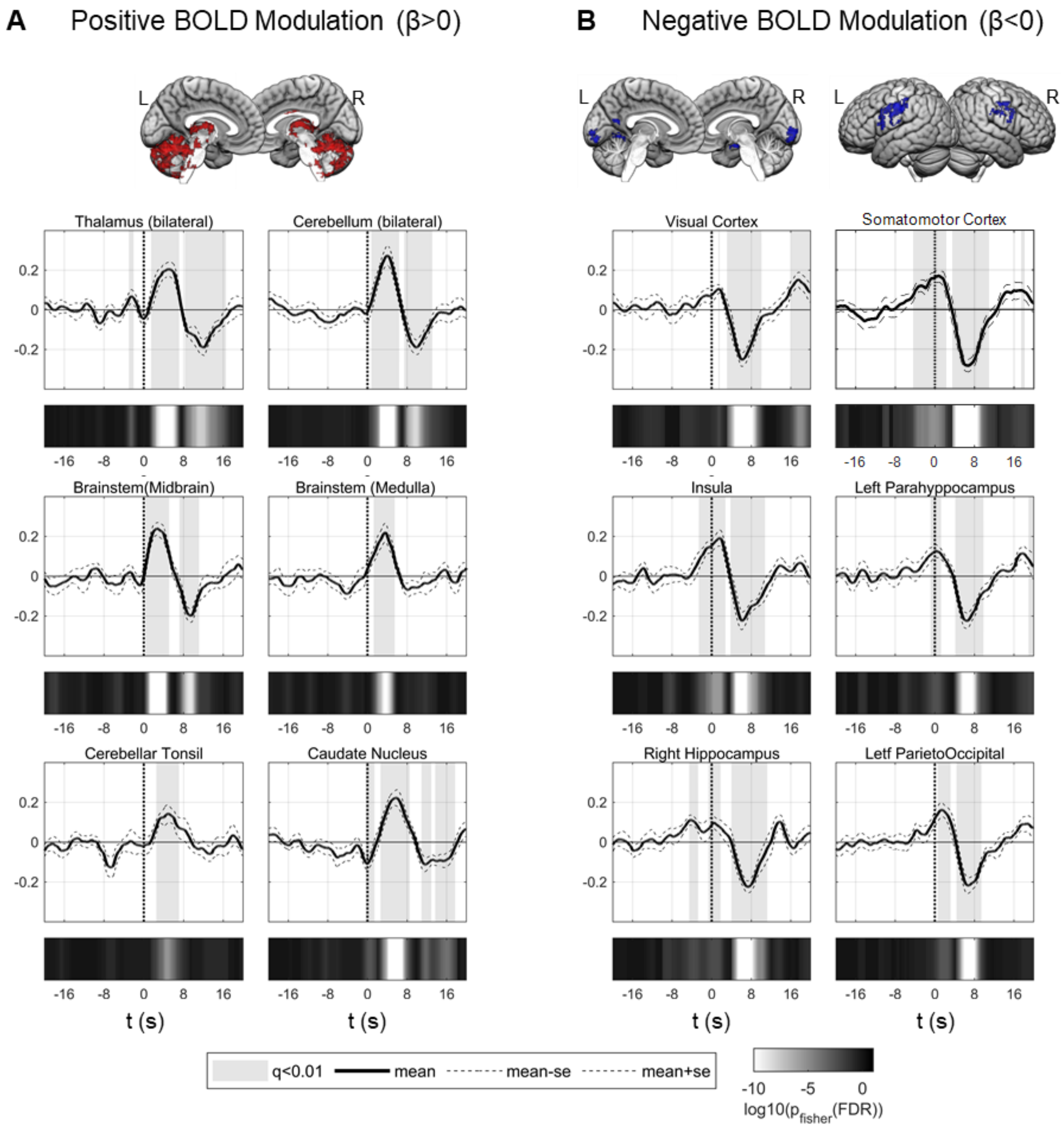


*Figure S5. Correlation between slow-wave amplitude and the magnitude of BOLD-signal variations. Point-by-point correlation between slow-wave amplitude and BOLD-signal changes for positively (subcortical; A) and negatively (cortical; B) modulated brain areas identified from the regression analysis. The gray shadowed areas indicate significant effects at the group level after FDR correction for multiple comparisons ($q < 0.01$). Time $t = 0$ s corresponds to slow-wave onset (positive-to-negative zero crossing).*

**Figure S6**

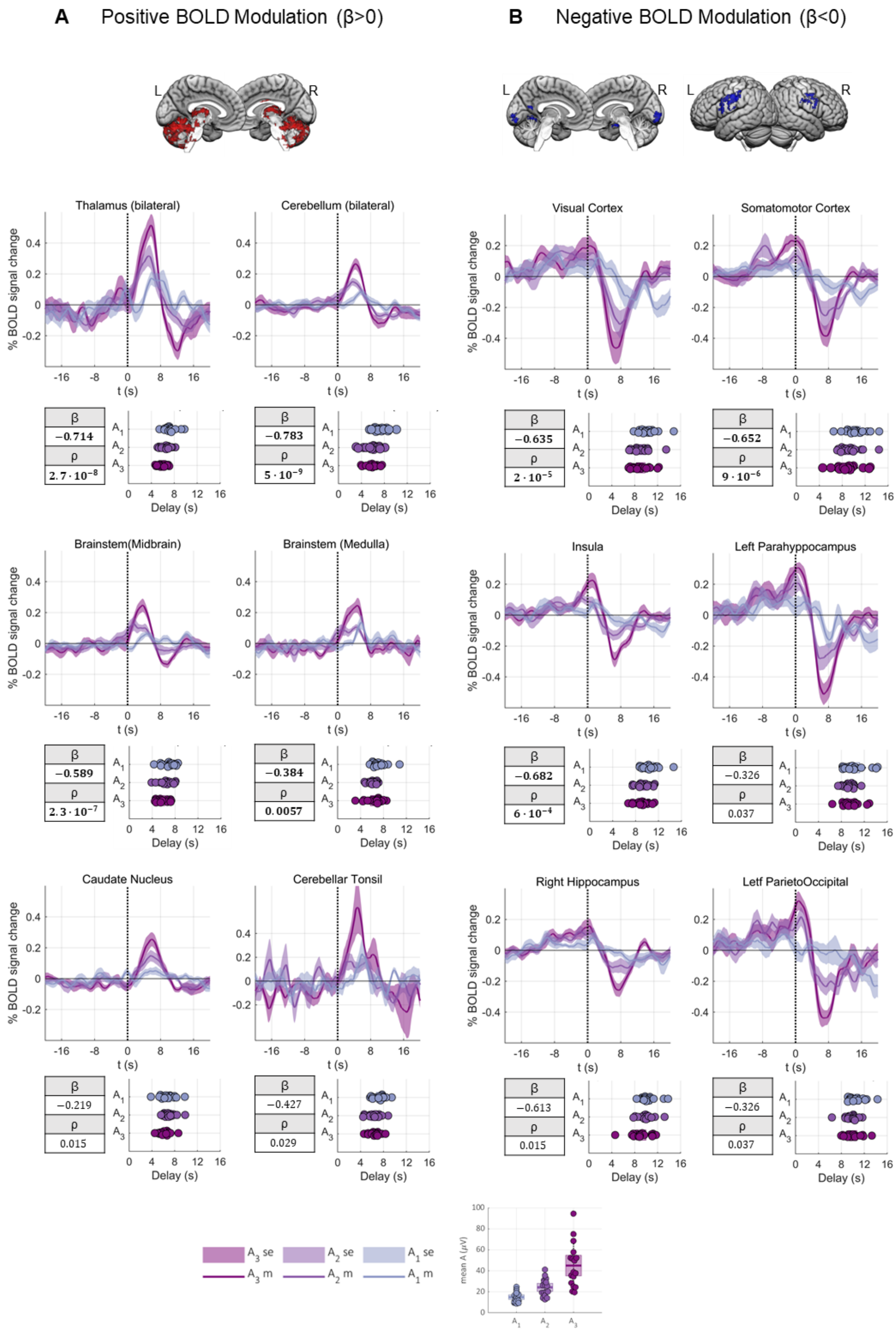


*Figure S6. Correlation between slow-wave amplitude and the latency of BOLD-signal variations. For each brain structure, the top panel shows the mean BOLD-signals (and relative standard errors) for three slow-wave classes defined based on amplitude percentiles ($A_1$: 0-33; $A_2$: 33-66; $A_3$: 66-100). The bottom panels show the relative delay of the positive (for subcortical structures; A) or the negative (for cortical areas; B) peaks in the BOLD-signal averaged for each subject within each slow-wave class. A regression analysis was performed to determine whether a relationship existed between slow-wave amplitude and the relative delays of hemodynamic peaks. Correspondent beta and p-values are reported in boxes for each region.*
